# Distinct life histories impact dikaryotic genome evolution in the rust fungus *Puccinia striiformis* causing stripe rust in wheat

**DOI:** 10.1101/859728

**Authors:** Benjamin Schwessinger, Yan-Jun Chen, Richard Tien, Josef Korbinian Vogt, Jana Sperschneider, Ramawatar Nagar, Mark McMullan, Thomas Sicheritz-Ponten, Chris K. Sørensen, Mogens Støvring Hovmøller, John P. Rathjen, Annemarie Fejer Justesen

## Abstract

Stripe rust of wheat, caused by the obligate biotrophic fungus *Puccinia striiformis* f. sp. *tritici*, is a major threat to wheat production world-wide with an estimated yearly loss of US $1 billion. The recent advances in long-read sequencing technologies and tailored-assembly algorithms enabled us to disentangle the two haploid genomes of *Pst*. This provides us with haplotype-specific information at a whole-genome level. Exploiting this novel information, we perform whole genome comparative genomics of two *P*. *striiformis* f. sp*. tritici* isolates with contrasting life histories. We compare one isolate of the old European lineage (PstS0), which has been asexual for over 50 years, and a Warrior isolate (PstS7 lineage) from a novel incursion into Europe in 2011 from a sexual population in the Himalayan region. This comparison provides evidence that long-term asexual evolution leads to genome expansion, accumulation of transposable elements, and increased heterozygosity at the single nucleotide, structural and allele levels. At the whole genome level, candidate effectors are not compartmentalized and do not exhibit reduced levels of synteny. Yet we were able to identify two subsets of candidate effector populations. About 70% of candidate effectors are invariant between the two isolates while 30% are hypervariable. The latter might be involved in host adaptation on wheat and explain the different phenotypes of the two isolates. Overall this detailed comparative analysis of two haplotype-aware assemblies of *P*. *striiformis* f. sp*. tritici* are the first steps in understanding the evolution of dikaryotic rust fungi at a whole genome level.

## Introduction

Fungi have complex life cycles involving clonal as well as sexual reproduction. Sex in fungi is often associated with dispersal, dormancy and survival under adverse conditions such as the absence of hosts (Anikster 1986). Sexual reproduction creates genotypic diversity and enables adaptation to new environments by combining beneficial alleles and eliminating harmful genetic traits (Seudre et al. 2018; Heitman 2015). The advantage of asexual reproduction, on the other hand, is the rapid multiplication and spread of the most well-adapted individuals. In some species, sexual reproduction is so rare or cryptic that the sexual cycle historically was thought to be absent and only recently has been discovered (Zeyl 2009; Taylor et al. 2015). Studies of the genomic architecture of fungal pathogens with different reproduction strategies can help us to understand the genetic and genomic processes behind pathogen adaptation. This knowledge is important for predicting the response of pathogens to challenges such as climate change, drugs, fungicides, host resistance or immunity, and will strengthen the development of methods for combating fungal pathogens (Gladieux et al. 2014).

Rust fungi, the *Pucciniales*, are a large group of fungal plant pathogens, some of which impact severely on agricultural and forest production (Lorrain et al. 2019). They depend on living host cells and have complex life cycles, which may comprise many spore types over two phylogenetically distinct host plants for completion of the sexual life cycle. *Puccinia striiformis* causes yellow (stripe) rust on cereals and grasses and has historically been restricted to temperate-cool and wet climates (Rapilly 1979). Since 2000, new aggressive strains adapted to higher temperatures have caused the expansion of the disease into warmer regions, and it is currently one of the most prevalent and damaging diseases on common wheat (*Triticum aestivum* L.) worldwide (Beddow et al. 2015; Wellings 2011; Savary et al. 2019; Singh et al. 2016). During the asexual life cycle, which recurs several times during the wheat growing season, *P. striiformis* produces dikaryotic urediniospores which reinfect wheat (Schwessinger 2017). These spores are wind borne and may spread over long distances to introduce exotic strains into new areas (Hovmøller et al. 2016; Ali et al. 2014). Historically, the sexual cycle of *P. striiformis* has been a great mystery and many years of search for the alternate host was fruitless until it was shown that barberry (*Berberis* spp.) can serve as the alternate host for completion of the sexual cycle (Jin et al. 2010). The significance of *Berberis* spp. as an alternate host for *P. striiformis* in nature is still unknown, but high levels of genetic diversity in *P. striiformis* populations in China and Pakistan indicate that sexual reproduction is taking place under natural conditions, and the near-Himalayan region is the likely center of genetic diversity for *P. striiformis* (Mboup et al. 2012; Duan et al. 2010; Ali et al. 2016; Ali et al. 2014). In contrast, reproduction of *P. striiformis* in Europe, America and Australia is clonal, and adaptation to host resistance happens through stepwise mutations (Chen et al. 2014; Hovmøller & Justesen 2007; Steele et al. 2001; Markell & Milus 2008). These mutations lead to the loss of recognition of *P. striiformis* effectors which previously were recognized by specific plant resistance (*R*) gene products and *P. striiformis* is then termed virulent to the specific *R* gene. In the absence of recognition, e.g. in a wheat plant that lacks the corresponding *R* gene, effectors suppress plant immune responses and facilitate the infection process (Ellis et al. 2014).

Only few clonal lineages of *P. striiformis* are responsible for the yellow rust epidemics worldwide, and stepwise mutation and clonal reproduction has been the major driving force for the evolution of new virulent races that can overcome widely deployed resistance genes (Walter et al. 2016; Hovmøller et al. 2008; Ali et al. 2017). Combination of virulence alleles by sexual or somatic recombination may also generate strains with new virulence profiles, the later was recently demonstrated for the *Puccinia graminis* f. sp. *tritici* isolate Ug99 (Li et al. 2019). The exact frequency of these mechanisms in nature is not yet known, but it may be more common than previously anticipated (Feng Li et al. 2019). Based on observational studies of more than 2000 alive isolates representing world-wide collections of *P. striiformis*, we have noted that isolates linked to long-term asexual lineages generally produce fewer teliospores compared to isolates from areas with sexual populations, and compared to first generation progeny isolates raised from sexual reproduction on *B. vulgaris* in particular (Rodriguez-Algaba et al. 2014). This observation is supported by Ali et al. 2010, 2014, where isolates from China, Pakistan and Nepal, which is considered center of genetic diversity for *P. striiformis*, produced more telia than isolates from e.g., Europe (Ali et al. 2010; Ali et al. 2014). Teliospores are obligatory for completing the sexual life cycle and this indicates that these long-term clonal lineages may have a reduced ability to undergo sexual reproduction (Ali et al. 2010).

In 2011, a new *P. striiformis* lineage, PstS7, known generally as Warrior was detected in high frequency on both wheat and triticale (*x Triticosecale*) in many European countries (Hubbard et al. 2015; Hovmøller et al. 2016). It was able to cause disease on cultivated host varieties that had previously been resistant (Sørensen et al. 2014). Moreover, it produced high amounts of teliospores and was able to complete the sexual cycle on barberry under experimental conditions (Rodriguez-Algaba et al. 2014). Warrior is genetically very different from the clonal lineages present in Europe before 2011 (e.g., PstS0) and similar to isolates originating from the centre of diversity in the near-Himalayan region (Hovmøller et al. 2016; Hubbard et al. 2015; Ali et al. 2014). Determining the mechanisms behind the generation of new virulent strains of *P. striiformis* such as Warrior is important. However, sequencing technologies available to date have provided only limited understanding of genetic recombination, comprising somatic, parasexual and sexual recombination, as a driving force for generating new strains of the pathogen.

Until recently, most draft genome assemblies of *P. striiformis* were based largely on short-read sequencing data (Cantu et al. 2011, 2013; Zheng et al. 2013). However, short-read sequencing data is not well suited to the dikaryotic rusts due to the high proportion of repetitive sequences (>50%), high heterozygosity, leading to the difficulty in phasing and high level of fragmentation of the two haploid genomes. Hence, short-read assemblies typically lack contiguity and individual haplotype information. Single molecule sequencing technologies such as PacBio and Nanopore produce far longer reads, which in combination with novel assembly algorithms have provided more accurate (Xia et al. 2018) and complete genomes with effective haplotype phasing (Schwessinger et al. 2018). Here we provide a genome assembly of a Warrior isolate sampled in Denmark (Pst-DK0911) using long-read sequencing and a diploid-aware genome assembler (Schwessinger et al. 2018; Chin et al. 2016). We previously produced a highly phased and contiguous assembly of Pst-104E (Schwessinger et al. 2018), an Australian isolate, which represents the first pathotype of *P. striiformis* detected in Australia in 1979 (Wellings & McIntosch 1990). This isolate is almost identical to European isolates belonging to the PstS0 lineage with respect to virulence and genotype (Hovmøller et al. 2008; Thach et al. 2016; Steele et al. 2001). The PstS0 lineage is strictly clonal (Hovmøller et al. 2002) and evolution of new races by stepwise mutations can be traced back to at least 1950s (Hovmøller & Justesen 2007; Thach et al. 2016). Isolates of the PstS0 lineage produce less teliospores with reduced ability to germinate and to complete the sexual life cycle on *Berberis* spp. (Ali et al. 2010).

This new genome assembly allows whole genome comparisons between two isolates representing contrasting lifestyles of *P. striiformis.* It provides detailed insights into the adaptive divergence and evolution of *P. striiformis* and rust fungi in general. Specifically, we explore questions around the impact of sexual and asexual reproduction on structural genomic evolution, the generation of genetic diversity, and host adaptation.

## Methods and materials

### *Puccinia striiformis* f. sp. *tritici* pathotype, growth conditions and spore amplification

The Pst-DK0911 isolate was purified from a single lesion sampled from the commercial wheat cv. Holeby in Denmark in 2011. The isolate was race phenotyped according to Hovmøller et al. 2017 and genotyped as described previously (Hovmøller et al. 2016; Rodriguez-Algaba et al. 2014). The Pst-104E is an isolate of the pathotype 104E137A- and was collected from the field in 1982 (Plant Breeding Institute accession number 821559=415). The sequencing and analysis of this isolate was described previously (Wellings 2007; Schwessinger et al. 2018). The same isolate ID was used in the study of Hovmøller et al. (2008), confirming the immediate connection between the old NW-European population and the Australian *P. striiformis* population up to 2001. The Pst-DK0911 isolate was multiplied on susceptible seedlings of wheat cv. Morocco, which were inoculated using the method described (Sørensen et al. 2016). The inoculated seedlings were misted with water and incubated at 10°C in the dark for 24 h and 100% relative humidity (dew formation) in an incubation room. After incubation the inoculated plants were transferred to quarantine spore-proof greenhouse cabinets with a temperature regime of 17°C day and 12°C night, and a light regime of 16 h photoperiod of natural light and supplemental sodium light (100 μmol s^−1^ m^−2^), and 8 h dark. The plants were covered with cellophane bags 5-7 days after infection (dai) to avoid cross-contamination before sporulation. The urediniospores were harvested at 15–20 dai and freeze-dried before DNA extraction.

### DNA extraction and genome sequencing

We extracted DNA from freeze-dried urediniospores as described previously (Schwessinger & Rathjen 2017) with few modifications. A volume of 875 μl lysis buffer was used per 60 mg spores in a 2 ml eppendorf tube. Library preparation and PacBio sequencing were performed at the Earlham Institute (Norwich, UK). One DNA library was sequenced on a PacBio RSII instrument using P6-C4 chemistry and 16 SMRT cells were sequenced in total. Moreover, a DNA sample from the same isolate was sequenced using Illumina short read technology. A TruSeq library was sequenced on a HiSeq 2000 instrument as a 100 bases paired-end library at the Danish National High-Throughput DNA Sequencing Centre (Copenhagen, Denmark).

### RNAseq

Pst-DK0911 was point inoculated on leaves of the susceptible cv. Avocet S with a pipette using 5 µl spore suspension in 3M^TM^ Novec^TM^ 7100 (2 mg spores per ml) as described previously (Sørensen et al. 2016) without fixation of leaves. Three replicates were prepared and leaf segments of 10-12 mm of the 2^nd^ leaves of each replicate were harvested at 7 and 9 dai for transcriptome sequencing. We isolated total RNAs with the RNeasy Plant Mini kit (Qiagen) from 100 mg of infected leaf tissues including a DNaseI (Qiagen) treatment according to the manufacturer’s instructions. Transcriptome libraries were generated using the TruSeq Stranded Total RNA with Ribo-Zero plant kit (Illumina) according to the standard protocol to remove cytoplasmic, mitochondrial and chloroplast ribosomal RNA. Next-generation sequencing (NGS) for 6 multiplexed samples was performed using an Illumina HiSeq2000 sequencer at BGI (Copenhagen, Denmark).

### Genome Assembly

For genome assembly we used FALCON and FALCON -Unzip (Chin et al. 2016) github tag 1.7.4 with the parameters described in Supplemental Data 1 and 2. We checked the resulting contigs for eukaryotic and prokaryotic contamination by blastn searches against the NCBI nucleotide reference database (downloaded 04052016) (Altschul et al. 1990). None of the contigs showed signs of contamination as estimated via best blast hits at a given position. We performed two manual curation steps. In the first step we reasoned that some of the primary contigs without corresponding haplotigs might actually represent haplotigs that could not be assigned to their respective primary contigs in the assembly graph because of too large a difference between the two haplotypes. We aligned all primary contigs without haplotigs to primary contigs with haplotigs using MUMmer version 3 (Kurtz et al. 2004). We screened the best alignments of each primary contig without haplotig for percentage alignment, length of alignment, and if they aligned to regions in the primary contigs that previously had not been covered by a haplotig alignment. Using this approach, we re-assigned five primary contigs without haplotigs (∼6 Mb) to haplotigs.

### Repeat annotation and analysis

Repeat regions were annotated as described previously (Schwessinger et al. 2018) using the REPET pipeline v2.5 (Quesneville et al. 2005; Flutre et al. 2011; Maumus & Quesneville 2014) for repeat annotation in combination with Repbase v21.05 (Bao et al. 2015). For de novo identification, we predicted repeat regions in primary contigs of Pst_104E_v13 and DK0911_v04 independently using TEdenovo. We combined the resulting de novo repeat libraries with the script ‘RemoveRedundancyBasedOnCI.py’ within the REPET pipeline using the configurations provided in Supplemental Data 3. We named the resulting repeat library 104E_p_DK0911_p_classif. We annotated the primary contigs and haplotigs of both assemblies separately using three repeat libraries (repbase2005_aaSeq, repbase2005_ntSeq, and 104E_p_DK0911_p_classif). We generated superfamily classifications as described previously (Schwessinger et al. 2018). See jupyter notebooks DK0911_TE_filtering_and_summary_p_contigs_fix.ipynb, DK0911_TE_filtering_and_summary_h_contigs_fix.ipynb, Pst_104E_v14_TE_filtering_and_summary_p_contigs_fix.ipynb, and Pst_104E_v13_TE_filtering_and_summary_h_contigs_fix.ipynb in the github repo for full analysis details.

We analyzed the relative identity distribution of TEs as proxy of TE age as previously described (Maumus & Quesneville 2014). The average identity and genome coverage per TE family is provided by the REPET pipeline. We used these tables as input and plotted the relative coverage frequency of TE families within average identity size bins of one (Supplemental Figure 1A). We plotted the cumulative difference in TE genome coverage between Pst-104E and Pst-DK0911 against the average TE family identity normalizing the current observed TE coverage difference to one. See jupyter notebook DK0911_TE_variation_analysis.ipynb for details.

### Gene model annotation

We annotated genes on primary contigs and haplotigs independently. We combined RNAseq guided *ab initio* prediction using CodingQuarry v2.0 (Testa et al. 2015) and braker v1.9 (Hoff et al. 2016) with *de novo* transcriptome assembly approaches using Trinity v2.2.0 (Grabherr et al. 2011) and pasa v2.0.1 (Haas et al. 2008). Gene models were unified using evidencemodeler v1.1.1 (Haas et al. 2008) using the weights given in Supplemental Data 4.

We mapped the trimmed RNAseq reads described in this study (see above) and previously (Dobon et al. 2016; Schwessinger et al. 2018) against the complete assembly (primary contigs and haplotigs) using hisat2 v2.1.0 (--max-intronlen 10000 --min-intronlen 20 --dta-cufflinks) (Kim et al. 2015). For *ab initio* predictions we reconstructed transcripts using stringtie v1.3.3b (-f 0.2) (Pertea et al. 2015). We ran CodingQuarry (-d) in the pathogen mode using SignalP4 (Petersen et al. 2011) for secretome prediction on the soft-masked genome using RepeatMasker v4.0.5 (-xsmall -s -GC 43). Similarly, we used the stringtie reconstructed transcripts as training set for the *ab initio* prediction pipeline braker 1 v1.9 (Hoff et al. 2016) and used the non-repeat masked genome as reference.

We used Trinity v2.2.0 to obtain *P*. *striiformis* f. sp*. tritici* transcripts both in the *de novo* mode and the genome guided mode. All RNAseq samples contained host and pathogen RNA as they were prepared from infected wheat tissue. We first mapped all reads to the genome using hisat2 (see above). We extracted mapped RNAseq reads using picard tools SamToFastq. Only those reads that mapped against *P*. *striiformis* f. sp*. tritici* contigs were used in the *de novo* pipeline of Trinity (--seqType fq). For genome guided assembly we used bam files generated with hisat2 as starting point for Trinity (--jacard_clip, --genome_gudied_max_intron 10000). We used the pasa pipeline v2.0.2 to align both sets of Trinity transcripts against *P*. *striiformis* f. sp*. tritici* contigs with blat and gmap.

The different gene models were combined using evidencemodeler v.1.1.1 to get the initial gene sets for primary contigs and haplotigs. These were filtered for homology with proteins encoded in transposable elements. We used blastp to search for homology in the Repbase v21.07 peptides database with an e-value cut-off of 1*e^-10^. In addition, we used transposonPSI (Bao et al. 2015). We used the outer union of both approaches to remove genes coding for proteins associated with transposable elements from our list of gene models.

### Protein annotation

We performed protein annotation as described previously including homology to swissprot proteins (uniref90, downloaded 22/09/2016) (Bairoch & Apweiler 2000), to carbohydrate-active enzymes (dbCAN, downloaded 22/09/2016) (Yin et al. 2012), to peptidases (MEROPS v10.0) (Rawlings et al. 2016) (see jupyter notebooks in the protein_annotation folder on github). We complemented this analysis by interproscan v5.21-60 (-iprlookup -goterms -pa) (Jones et al. 2014), eggnog-mapper v0.99.2 (-m diamond and –d euk) (Huerta-Cepas et al. 2016), SignalP 3 (Bendtsen et al. 2004), and EffectorP v2.0 (Sperschneider et al. 2015, 2018).

In case of Pst-DK0911 proteins labelled as “candidate effectors” or “EffectorP” belong to the same protein group. They are defined as secreted proteins using SignalP3 followed by being labelled as “candidate effectors”/”effectorP” by the machine learning program EffectorP v2.0.

In case of Pst-104E protein grouping was described in detail previously (Schwessinger et al. 2018). Briefly, “effectorP” labelled proteins were defined as above except for using EffectorP v1.0. “Candidate effectors” are the outer union of “effectorP” labelled proteins and secreted proteins that are upregulated in haustoria or during the plant infection process when compared to spore stages.

### BUSCO analysis

We used BUSCO3 to identify core conserved genes and to assess genome completeness (Simão et al. 2015). In all cases we ran BUSCO3 in the protein mode using the basidiomycota reference database downloaded 01/09/2016 (-l basidiomycota_odb9 -m protein).

### Inter-haplotype variation analysis

We performed the inter-haplotype variation analysis similar as described previously (Schwessinger et al. 2018). Trimmed Illumina reads derived from DK0911, Pst_104E, and PST78 were mapped against the genomes (DK0911 and Pst_104E) using BWA-MEM v0.7.15-r1142-dirty using the standard parameters (Li 2013). We called SNPs in bulk using freebays (Garrison & Marth 2012) and following hard filtering by vcffilter with DP > 10 and QUAL > 20 (vcflib, 2017b). Variants for DK0911, Pst_104E, and PST78 resequencing data were parsed out and summarized with real time genomic vcfstats v3.8.4 (rtg-tool, 2017a).

We performed large-scale variation analysis as described previously using nucmer and assemblytics (Kurtz et al. 2004; Nattestad & Schatz 2016). See jupyter notebooks DK_0911_nucmer_assemblytics_mapping.ipynb, DK_0911_assemblytics_analysis.ipynb, and DK_0911_Pst104E_assemblytics_plot.ipynb for details.

### Within genome allele variation analysis

We used proteinortho v5.16 in synteny mode with default parameters (-synteny) to identify alleles between the primary assembly and haplotigs (Lechner et al. 2011). We restricted our analysis to high quality allele pairs that are located on linked primary contigs and haplotigs. We assessed the variation of high-quality allele pairs by calculating the Levenshtein distance on the codon-based CDS alignments. The pairwise protein alignments were generated with muscle v3.8.31 (Edgar 2004), and converted into codon-based CDS alignments using PAL2NAL v14 (Suyama et al. 2006). Analysis details can be found in jupyter notebook DK0911_vs_Pst104E_gene_pair_analysis.ipynb.

### Coverage analysis and identification of unphased regions in primary contigs

We identified unphased regions in Pst-DK0911 as reported previously for Pst-104E (Schwessinger et al. 2018). We performed detailed read depth coverage analysis to obtain a genome level insight into the relative quantity of fully phased, homozygous collapsed, and hemizygous regions in the two *P*. *striiformis* f. sp*. tritici* genomes. We determined the sequencing depth in 1-kb sliding windows with 200-base intervals. Read depth coverage was calculated by mapping short reads derived from each *P*. *striiformis* f. sp*. tritici* isolate against its own genome using BWA-MEM v0.7.15-r1142-dirty and the standard parameters (Li 2013). We mapped reads against primary contigs only (p) and both primary contigs and haplotigs (ph). We normalized sequencing depth to one at the main “haploid” sequencing depth peak on primary contigs when mapping against primary contigs and haplotigs. In a perfect fully phased assembly this is the main sequencing depth peak, because of the absence of any homozygous collapsed regions. We plotted the normalized sequencing depth for primary contigs (both p and ph mapping), haplotigs (ph mapping), regions in primary contigs that have corresponding phased haplotigs and regions in primary contigs without assigned haplotigs. In addition, we used publicly available long-read genomes and corresponding Illumina short-reads for three additional *P. striiformis* isolates. This includes *P. striiformis* f. sp. *tritici* 93-210 (SAMN08200485), *P. striiformis* f. sp. *hordei* 93TX-2 (SAMN08200486), and *P. striiformis* 11-281 (SBIN00000000) (Li et al. 2019; Xia et al. 2018). See Supplemental Data 5 for SRA identifiers. For details on analysis specifics see jupyter notebook DK0911_SRM_cov_DK0911_on_DK0911.ipynb, Pst_104E_SRM_cov_Pst_104E_on_Pst_104E.ipynb, DK0911_Pst104E_comperative_coverage_analysis.ipynb, and 20200129_het_cov_revision_v01.ipynb.

### Short-read k-mer frequency analysis to estimate levels of homo- and heterozygosity

We investigated levels of homo- and heterozygosity directly from short-read sequencing data for several additional *P. striiformis* f. sp. *tritici* isolates linked to recent incursions from the Himalayan region plus appropriate controls (Li et al. 2019; Xia et al. 2018; Schwessinger et al. 2018; Bueno-Sancho et al. 2017; Hubbard et al. 2015). We performed k-mer analysis on raw Illumina short-read using jellyfish v2.2.6. We first invoked the count command with the k-mer size of 21 (-m 21) and size of 1000000000 (-s 1000000000). We converted the resulting count file into a histogram using the histo command of jellyfish (Marçais & Kingsford 2011). We plotted k-mer frequency spectrums and estimated heterozygosity using GenomeScope2 online last accessed 2020/02/12 (Ranallo-Benavidez et al. 2019). See Supplemental Data 5 and 20200129_genome_size_revision_v01.ipynb for details.

### Cross isolate genome wide presence-absence analysis

We aimed at determining presence absence polymorphism at the whole genome level. We first mapped short reads derived from one *P*. *striiformis* f. sp*. tritici* isolate (‘query’) against the complete genome (ph) of the other isolate (‘subject’) using BWA-MEM v0.7.15-r1142-dirty using standard parameters. We then calculated sequencing depths in 1 kb sliding windows as described above. We defined regions that fell below 10% normalized sequencing depth in the cross mapping but not self mapping experiments as absent in the ‘query’ isolate. We used bedtools (Dale et al. 2011; Quinlan & Hall 2010) to identify genes which are completely contained within these regions and defined them as ‘variable genes’. We estimated the distribution of the expected number of variable genes using a permutation test randomly reshuffling the absent regions five thousand times. We used this random distribution to calculate a p-value testing the hypothesis if the observed number of variable genes is different from the observed random distribution at a p-value cut-off of 0.05. We performed a similar analysis for TEs with the important difference to determine variable TE content at single base pair resolution and with two thousand five hundred permutations. In addition, we performed similar tests on the variable TE content using reciprocal whole genome alignments as input. We aligned each genome against each other with the mummer package v4. Initially, alignments with nucmer used -l 20 -c 65 --max-match. Alignments were filtered with delta-filter requiring no uniqueness - u 0 but a minimum sequence identify of 95% -i 95 (see notebook Mummer_DK0911_Pst_104E.ipynb and DK0911v04_Pst104E_presence_absence.ipynb for more detail). We converted mummer delta files into bed files for input in the presence absence analysis as described above. Details on this analysis can be found in jupyter notebooks DK0911_SRM_cov_Pst_104E_on_DK0911.ipynb, Pst_104E_SRM_cov_Pst_104E_on_Pst_104E.ipynb, Mummer_DK0911_Pst_104E.ipynb, and DK0911v04_Pst104E_presence_absence.ipynb.

### Ortholog analysis

We performed orthology analysis orthofinder/2.2.6 (Emms & Kelly 2018) of all non-redundant protein sets with publicly available *Pucciniomycotina* genomes. Protein sets were downloaded from MycoCosm (Grigoriev et al. 2014) (see Supplemental Data 6 for details). *Zymoseptoria tritici* and *Verticillium dahliae* protein sets were also included as out-group.

We defined singletons between the two isolates as proteins that were in orthogroups that did not contain a member from the other isolate. We performed enrichment analysis interrogating if different gene categories are enriched for singletons over others. For this we used Fisher’s exact test and bonferroni correction executed in python (McKinney 2010). Detailed analysis can be found in jupyter notebook DK0911_vs_Pst104E_synteny.ipynb, Pst104E_vs_DK0911_synteny.ipynb, DK0911_vs_Pst104E_reciprocal_gene_pair_analysis.ipynb, and DK0911v04_Pst104E_presence_absence_orthology.ipynb.

### Synteny analysis

We performed detailed synteny analysis to identify the best reciprocal gene pairs between the two isolates. In addition, we investigated if certain gene categories are more syntenic than others considering parameters like maximum synteny block size and number of allowed gaps within a synteny block.

We identified best reciprocal gene pairs as follows. For each orthogroup we identified the reciprocal gene pairs or groups belonging to each isolate. If there was more than one gene pair candidate we identified the pair(s) for which each downstream and upstream gene was conserved in terms of orthogroup identity as well. In the case of multiple candidate gene pairs we assigned gene pair status to pairs that formed the longest synteny block when allowing for a maximum synteny gap number of eight.

We identified synteny blocks using MCScanX (Wang et al. 2012) inspired by previous analysis (Zhao & Schranz 2019). We ran MCScanX (Wang et al. 2012) and Diamond (Buchfink et al. 2015) within the Synnet.sh script with following parameters. MCScanX was run fixing the -w flag to 0, the -s flag to 3, and varying the -m flag with the following values: 0, 1, 2, 3, 4, 5, 6, 7, 8, 9, 10, 15, 25, 35. These settings allow for the identification of synteny blocks involving at least three genes with a varying number of allowed gaps within the synteny block. See Supplemental Figure 2 for an illustration of gaps within synteny blocks. Diamond was run with the following flags: --max-hsps 1 -k 25 -f 6 -b 5. Resulting *.collinearity files were parsed and further analyzed as described above. Detailed description of downstream analysis can be found in in jupyter notebook DK0911_vs_Pst104E_reciprocal_gene_pair_analysis.ipynb and DK0911v04_Pst104E_presence_absence_orthology.ipynb.

### Gene pair variation analysis

We assessed the variation between the gene pairs of the two isolates using three approaches. We calculated the Levenshtein distance based on the codon based CDS alignments and on the protein alignments of each gene pair. We calculated the dN:dS ratios by using these two alignment sets with yn00 paml version 4.9 (Yang & Nielsen 2000). The protein sequences of each gene pair was aligned using muscle v3.8.31 (Edgar 2004), and codon-based alignments were generated using PAL2NAL v14 (Suyama et al. 2006). The Levenshtein distance was calculated in python using the distance module v0.1.3. Analysis details can be found in jupyter notebook DK0911_vs_Pst104E_reciprocal_gene_pair_analysis.ipynb.

### Telomere analysis

We estimated telomere length and telomere patterns using computel (Nersisyan & Arakelyan 2015) providing a haploid chromosome number of 18 (-nchr) and the genome size (-lgenome) of the primary assemblies. The haploid chromosome number is based on the observation that closely related rust fungi such as *P. graminis* f. sp. *tritici* (Boehm et al. 1992; Sperschneider et al. 2020; Li et al. 2019) and *Melampsora lini* (Boehm 1992) have 18 chromosomes haploid. Hence, it is the most parsimonious assumption that *P. striiformis* f. sp. *tritici* haploid chromosome number is 18.

### Genome architecture analysis

We analyzed various features between certain gene families and in regards to TEs using bedtools (Quinlan & Hall 2010) and pybedtools (Dale et al. 2011) as described previously (Schwessinger et al. 2018). See jupyter notebook DK0911_v04_effector_analysis.ipynb.

### Mitochondrial genome assembly, annotation, and comparison

To assemble the Pst-DK0911 mitochondrial genome, we extracted mitochondrial reads from whole genome PacBio reads of Pst-DK0911 by mapping them to the closely related *Phakopsora meibomiae* (soybean rust pathogen) mt-genome (Stone et al. 2010) using bwa version 0.7.15-r1140 (Li 2013). The reads were *de novo* assembled with Canu v1.6 (Koren et al. 2017). Canu produced a single circular contig of ∼124 kb. The circularity of the contig was confirmed by dot plot using Gepard v-1.40 (Krumsiek et al. 2007). This produced a typical dot plot with identical repeats at both ends, indicating a circular topology. The repeated sequences at the ends of the contig were removed and the remainder circularized using circlator v-1.4.0 (Hunt et al. 2015). The final circularized contig was of 101,813 bp. The contig was polished/error corrected with two iterations of long reads using PacBio’s GenomicConsensus Arrow (2019) and five iterations of Illumina short reads using pilon v-1.22 (Walker et al. 2014). The polished mt-genome was annotated with two independent mt-genome annotation tools, GeSeq web browser, https://chlorobox.mpimp-golm.mpg.de/geseq.html (Tillich et al. 2017) using the *Phakopsora* mt-genome as a reference and MITO web server using genetic code 4 http://mitos.bioinf.uni-leipzig.de/index.py (Bernt et al. 2013). Annotations using these web browsers were checked individually to confirm gene boundaries as well as intron-exon boundaries by aligning the predicted genes with their orthologs in closely-related fungal species. Annotated tRNA genes were further evaluated and compared using tRNAscan-SE v2.0.3 (Lowe & Eddy 1997) and ARAGORN v1.2.38 (Laslett & Canback 2004). Introns were annotated using RNAweasel (Lang et al. 2007). We compared our mitochondrial genome with three previously published mitochondrial genomes that have a similar length of about 100kb including *P. striiformis* f. sp. *tritici* 93-210 (CM009485), *P. striiformis* f. sp. *hordei* 93TX-2 (CM009486), and *P. striiformis* f. sp. *tritici* CY32 (MH891489) (Li et al. 2020; Xia et al. 2018). We performed whole genome alignments using LASTZ (Harris 2007) within Geneious Prime version 2020.0.5. We visually inspected gene annotation between Pst-DK0911 and Pst-CY32.

## Results

### A high quality partially haplotype phased genome for Pst-DK0911, a Warrior (PstS7) isolate

We aimed to generate a haplotype-phased genome of the isolate Pst-DK0911 representing the PstS7 lineage (Ali et al. 2017) generally known as “Warrior” (Hovmøller et al. 2016) using PacBio long-reads (Supplemental Data 7, Table 1) assembled with FALCON-Unzip (Chin et al. 2016), which is designed to output haplotype phase blocks (haplotigs) for regions where homologous chromosomes vary significantly. The primary genome assembly of Pst-DK0911 is highly contiguous with 94 contigs and a contig N50 of 1.54 Mb, whereas the haplotigs are far more fragmented consisting of 1176 contigs. The 94 primary contigs totalled 74 Mb in size and the 1176 haplotigs constituted a size of 52 Mb (Table 1). Here haplotigs represent genome regions where the two haploid genome copies are significantly different. The genome was annotated based on new and previously published expression data from several different developmental and infection stages of *P*. *striiformis* f. sp*. tritici* on wheat (Dobon et al. 2016; Schwessinger et al. 2018). The Pst-DK0911 genome appears to be complete as measured by the identification of benchmarking conserved single copy orthologs (BUSCOs, basidiomycota odb9) (Simão et al. 2015). We could identify complete BUSCO hits for 97% (1290/1335) of the benchmarking set with only 2% (25/1335) missing. In addition, the assembly has a low level of duplicated BUSCOs (< 5%) indicating that it is completely phased and that primary contigs do not contain extensive duplicated regions or unassigned haplotigs. This makes our Pst-DK0911 genome assembly high quality and comparable to previous long-read genome assemblies for *P*. *striiformis* f. sp*. tritici* (Schwessinger et al. 2018; Xia et al. 2018) and therefore suitable for in depth comparative genomics with the other available partial haplotype phase assembly of Pst-104E (Schwessinger et al. 2018). In total we annotated 15,070 genes on primary contigs and 10,870 genes on haplotigs in Pst-DK0911. The overall gene length (1550 bp on primary contigs and 1420 bp on haplotigs) and average number of introns per genes (3.3 and 3.1 for primary- and haplotigs, respectively) is similar between primary contigs and haplotigs and near identical to our previous long-read assembly of Pst-104E (Table 1) (Schwessinger et al. 2018).

**Table 1:**
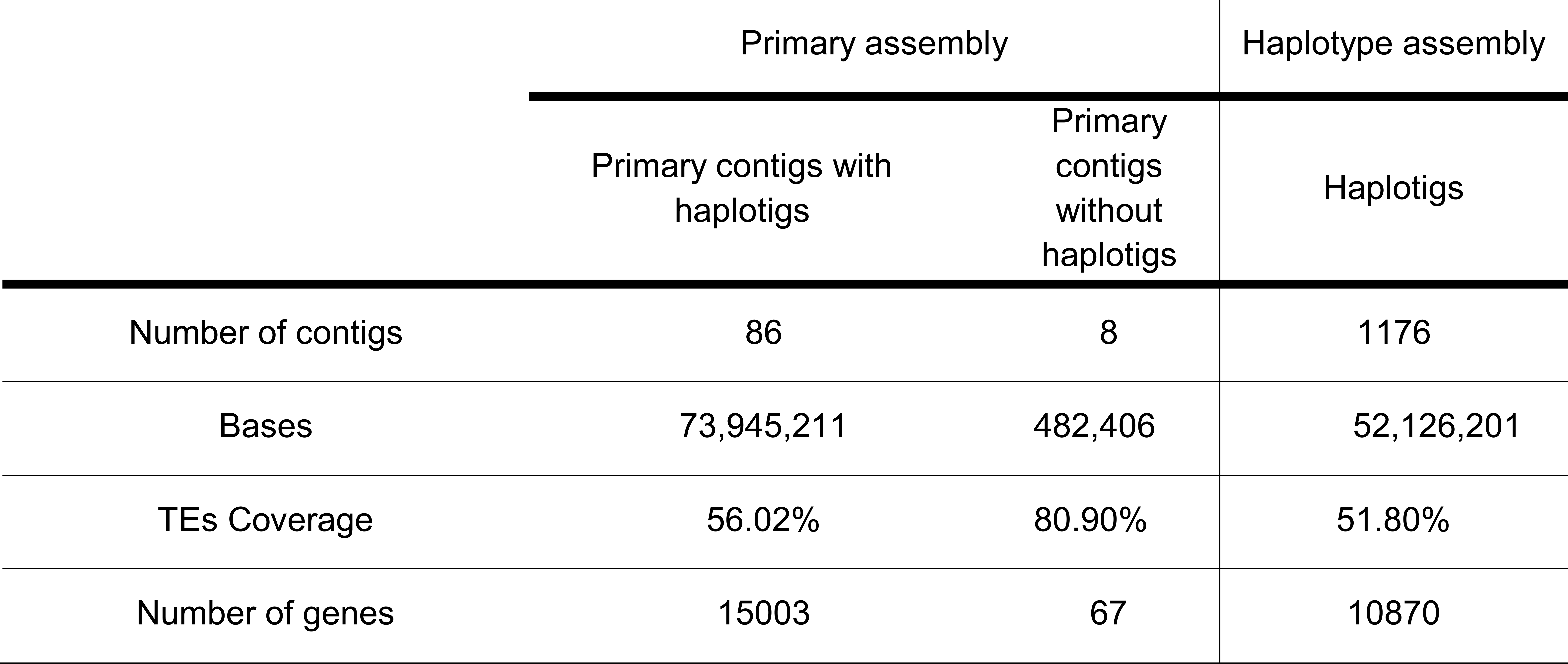
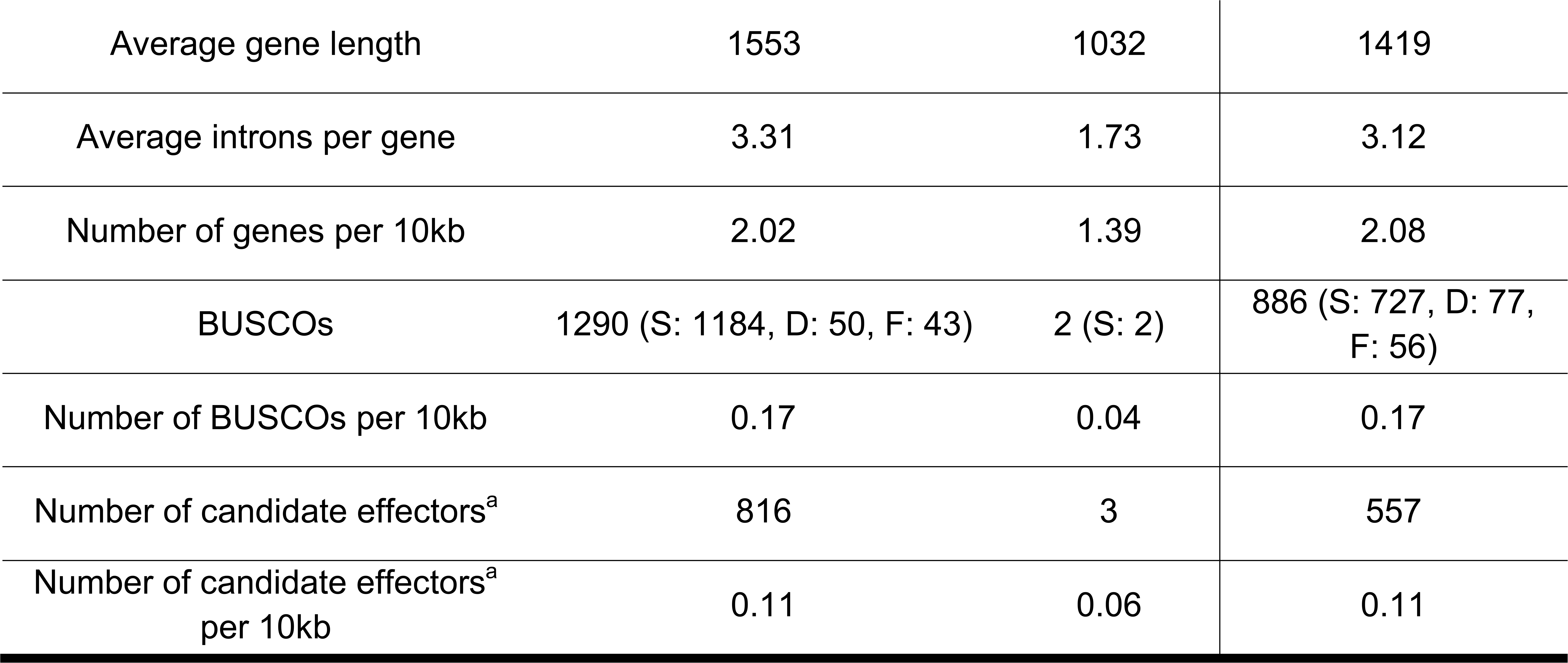
Summary of Pst-DK0911 genome assembly and annotation. Summary statistics for the genome assembly according to the three different contig categories as described in the main text. For the BUSCO annotation we used the following abbreviations. S single copy, D duplicated, F fragmented. ^a^Candidate effectors have been predicted based on the machine learning algorithm EffectorP2 only. In the BUSCO column the abbreviations are as follows: S single-copy, D duplicated, F fragmented

Potential molecular functions of all protein coding genes were assessed by searches against several databases (see Methods section). About 50% of the proteome contained at least one functional domain (Supplemental Data 7, Table 2). We predicted candidate effectors with the machine learning program EffectorP 2.0 (Sperschneider et al. 2018) applied to the secretome identified with SignalP 3.0 (Bendtsen et al. 2004). We identified 819 candidate effectors on primary contigs and 557 on haplotigs.

### A high-quality mitochondrial genome for Pst-DK0911

Using PacBio long-reads we were able to assemble a high-quality, circular mitochondrial genome for Pst-DK0911 with a length of 101,813 bp (Supplemental Figure 3), which displays a high structural and sequence conservation when compared to three previously published *P. striiformis* mitochondrial genomes (Li et al. 2020; Xia et al. 2018) (Supplemental Figure 4). The Pst-DK0911 mt-genome contains 14 protein coding genes that encode components of the electron transport chain, including three copies of the ATP synthase complex (*atp6*, *atp8* and *atp9*), seven subunits of the nicotinamide adenine dinucleotide ubiquinone oxidoreductase complex (*nad1*, *nad2*, *nad3*, *nad4*, *nad4L*, *nad5* and *nad6*), one cytochrome b (*cob*), and three subunits of cytochrome c oxidase (*cox1*, *cox2* and *cox3*). In addition, the mt-genome encodes a ribosomal protein (*rps3*) which is a frequent component of fungal mt-genomes (Korovesi et al. 2018; Losada et al. 2014), and a set of 24 tRNAs. All coding and non-coding genes are transcribed clockwise by the plus strand only. The protein coding regions are separated by large (800-2,100 bp) regions containing abundant AT-rich repeats.

### The difference in genome size between Pst-DK0911 and Pst-104E, representing two different *P*. *striiformis* f. sp*. tritici* lineages, is explained by the difference in TE content and suggests a recent TE burst

The overall genome composition and coding capacity (Table 1) of Pst-DK0911 is very similar to that of previous long-read assemblies (Schwessinger et al. 2018; Xia et al. 2018). At the same time the cumulative size of the primary contigs of Pst-DK0911 is 74 Mb whereas it is 83 Mb for Pst-104E and 85Mb for Pst-93-210. We investigated if this difference might be linked to differences in transposable element (TE) content. When analysing repetitive elements, approximately 56% of the Pst-DK0911 genome was annotated as encoding TEs, shared equally between Class I and Class II elements (Supplemental Figure 2). In the case of Class I retrotransposons, LTR Gypsy and Copia elements were by far the most abundant superfamilies. TEs belonging to the TIR order were the most abundant Class II DNA transposons. The overall distribution of TEs into superfamiles and their relative percentage identity compared to the family consensus sequence in Pst-DK0911 was very similar to what have previously been observed in Pst-104E with the most recent TE amplification peak centering around 90% family level sequence identity (Schwessinger et al. 2018) (Supplemental Figure 1A and 5). Here sequence conservation (identity) can be used as a proxy for TE age (Maumus & Quesneville 2014), so more recent TE duplication events display a greater sequence conservation than older TE copies, indicating that the expansion of TEs in Pst-104E and Pst-DK0911 has happened in recent history.

Yet TE genome coverage was distinct between the two genomes contributing to the difference in primary contig size (approximation of genome size) between Pst-DK0911 (Table 1) and Pst-104E (Schwessinger et al. 2018). TE content of Pst-DK0911 was 41 Mb when compared to 47 Mb in Pst-104E, which accounts for a 14% increase in TE content and for 54% of the observed genome size difference between the two genomes. Indeed, there appears to be a specific expansion of TEs in Pst-104E that is absent in Pst-DK0911 (Supplemental Figure 1B). About 40% of the difference in TE coverage in Pst-104E relative to Pst-DK0911 is explained by an expansion of TEs with an average sequence identity relative to the family consensus sequence ranging from 82 to 95% (Supplemental Figure 1B). It is also the relatively young TE families that demonstrate the largest difference in coverage between Pst-104E and Pst-DK0911, these belong mainly to the Class I LTR Gypsy and unclassified Class II elements (Supplemental Figure 1C and D).

### Comparatively low heterozygosity levels in Pst-DK0911 suggest long-term asexual reproduction contributes positively to the accumulation of genetic variations

In addition to a substantially smaller genome size, the Pst-DK0911 genome was phased to a much lower level than Pst-104E, reaching levels of 70% phasing compared with over 90% for Pst-104E (Schwessinger et al. 2018). This suggests that Pst-DK0911 is less heterozygous than Pst-104E, because lower levels of heterozygosity prevent the phasing of haplotypes during the assembly process (Chin et al. 2016). Hence, we tested heterozygosity levels in five independent ways. First, we determined the number of heterozygous single nucleotide polymorphisms (SNPs) when mapping Illumina short reads of the same isolate against its primary contigs only. Here Pst-104E had about 7.1 SNPs per kb (approx. 0.71% variation) which is consistent with previous reports of isolates belonging to the PstS0 and PstS1/2 lineage, e.g. Pst 93-210 6.4 SNPs/kb, Pst-21 5.0 SNPs/kb, Pst-43 5.3 SNPs/kb, Pst-130 5.4 SNPs/kb, Pst-87/7 6.6 SNPs/kb, Pst-08/21 7.6 SNPs/kb, and Pst-78 6.0 SNPs/kb (Schwessinger et al. 2018; Xia et al. 2018; Cantu et al. 2013, 2011; Hubbard et al. 2015; Radhakrishnan et al. 2019; Bueno-Sancho et al. 2017; Cuomo et al. 2016; Zheng et al. 2013). In contrast Pst-DK0911 had only 1.6 SNPs per kb (approx. 0.16% variation), which is dramatically less and consistent with our observation of reduced phasing in the Pst-DK0911 genome assembly. Secondly, we made use of the partially phased assemblies to estimate larger scale variations. When comparing Pst-DK0911 with Pst-104E we consistently observed that the cumulative variation size in different size bins is always larger in Pst-104E (Figure 1). In total, the larger scale structural variants covered 5.10 Mb (approx. 6.39% variation) in Pst-104E and 2.00 Mb (approx. 2.66% variation) in Pst-DK0911. Third, we analyzed the normalized read mapping coverage of primary contigs and haplotigs to estimate the level of heterozygosity and of hemizygosity versus the level of collapsed regions in the two genome assemblies. We normalized the read mapping coverage to one (‘haploid coverage’) using completely phased regions of each genome as reference when mapping reads against both primary contigs and haplotigs (Figure 2 B and D). When mapping against the primary contigs only (Figure 2A), nearly all regions of the Pst-DK0911 genome displayed two times (‘diploid’) coverage, suggesting a high level of similarity between the two haplotypes, such that reads from one haplotype are able to map to the other haplotype. In contrast, Pst-104E also showed a peak at 1x coverage which suggests the presence of hemizygous regions in its genome. Indeed, when mapping reads against primary contigs and haplotigs, Pst-104E showed a single strong coverage peak for all regions analyzed (Figure 2 B-D) including regions in primary contigs that lack an alternate haplotig. This supports the conclusion that Pst-104E contains hemizygous regions in its genome that are significantly different between the two haploid nuclei. In contrast, nearly all genome regions in Pst-DK0911 which lack a corresponding haplotig display 2x ‘diploid’ coverage indicating that these regions are collapsed in this genome assembly and present in both haploid nuclei. We also made use of three additional available long-read primary genome assemblies of *P. striiformis*, Pst-93-210, Psh-93TX-2, and Ps-11-281 (Li et al. 2019; Xia et al. 2018) to investigate if high levels of hemizygous regions are common. Indeed, all three genome assemblies display significant levels of hemizygous regions similar to Pst-104E based on normalized read mapping coverage of primary contigs (Supplemental Figure 6). Fourth, we estimated heterozygosity rates based on k-mer frequency profile of Illumina short-read sequencing data. This approach revealed relatively high heterozygosity rates in Pst-CY32 previously (Zheng et al. 2013) and is exemplified by a double peak in the k-mer frequency profile (Figure 3). This approach enabled us to extend our analysis to four more *P. striiformis* f. sp. *tritici* isolates that have been recently introduced to western wheat growing areas from the Himalayan region (Bueno-Sancho et al. 2017; Hubbard et al. 2015). Overall all five isolates recently derived from the Himalayan region, Pst-DK0911, Pst-11/08, Pst-Kranich, Pst-12/83, and Pst-12/86, display a single k-mer frequency peak when compared to the other non-Himalayan region isolates including Pst-104E (Figure 3 and Supplemental Data 5). Fifth, we analyzed the nucleotide variation of high confidence gene pairs (alleles) within each assembly (primary contigs versus linked haplotigs). A total of 52% (7472/14321) of all possible high-quality gene pairs in Pst-104E displayed variation on the CDS level (Levenshtein distance greater than zero) compared to 40% (4344/10870) in Pst-DK0911. The distribution of nucleotide variation of high quality gene pairs varied significantly between Pst-104E and Pst-DK0911 (median: 0.0053 vs. 0.0012, Mann-Whitney U test, p-value = 3.13e-93) with the first quartiles being more extended in Pst-104E compared to Pst-DK0911, while following quantiles are more extended in Pst-DK0911 (Figure 4). This suggests two different potential sources of gene pair variation. Overall Pst-104E is clearly more heterozygous than Pst-DK0911 in its genome architecture.

**Figure 1:**
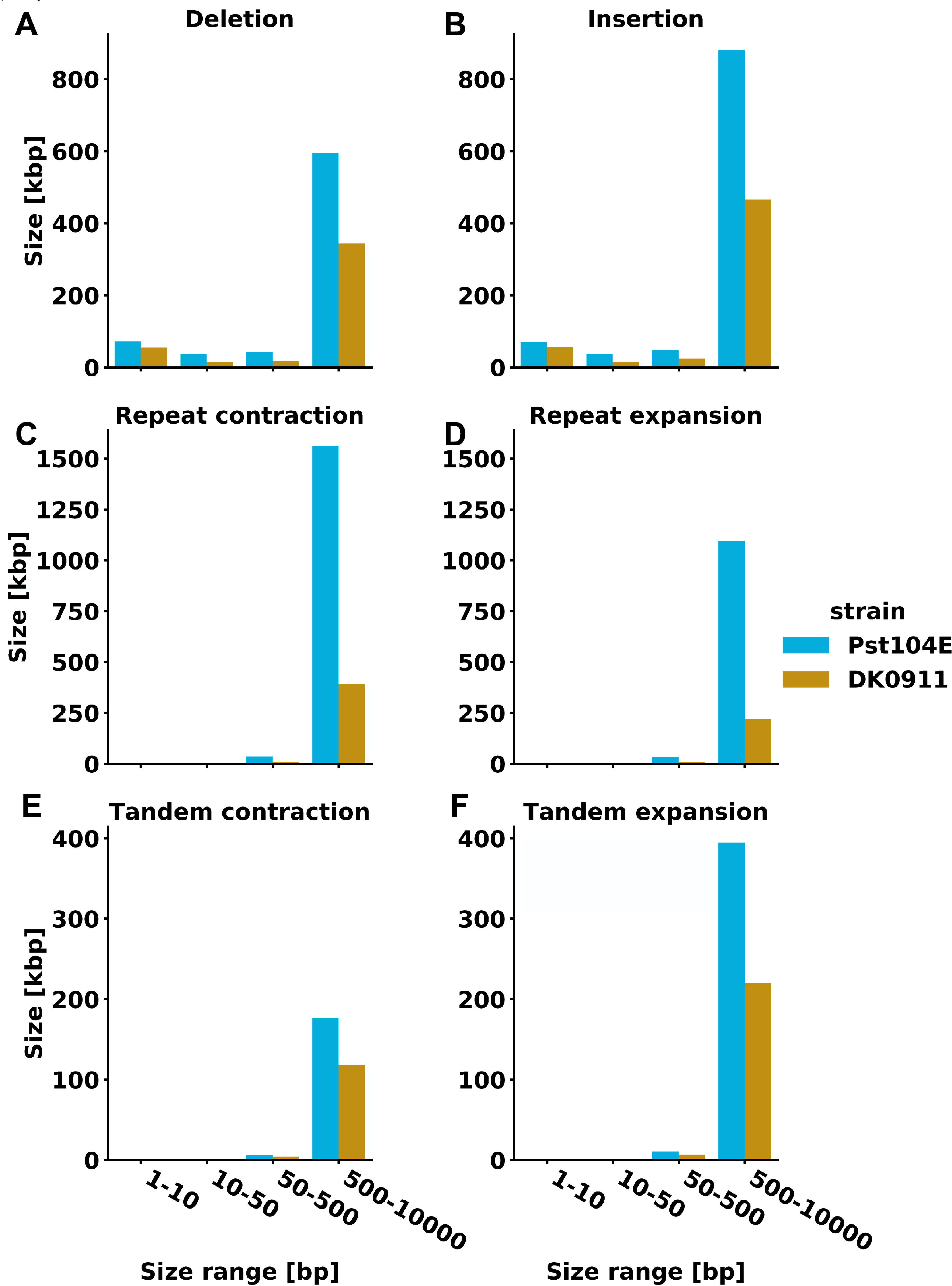
Reduced inter haplotype structural variations in Pst-DK0911. Summary of interhaplotype variation between primary contigs and their respective haplotigs, analyzed using Assemblytics. Each plot indicates the number of bases that are spanned by the specific variation category. Pst-DK0911 displays less variation then Pst-104E in all categories.

**Figure 2:**
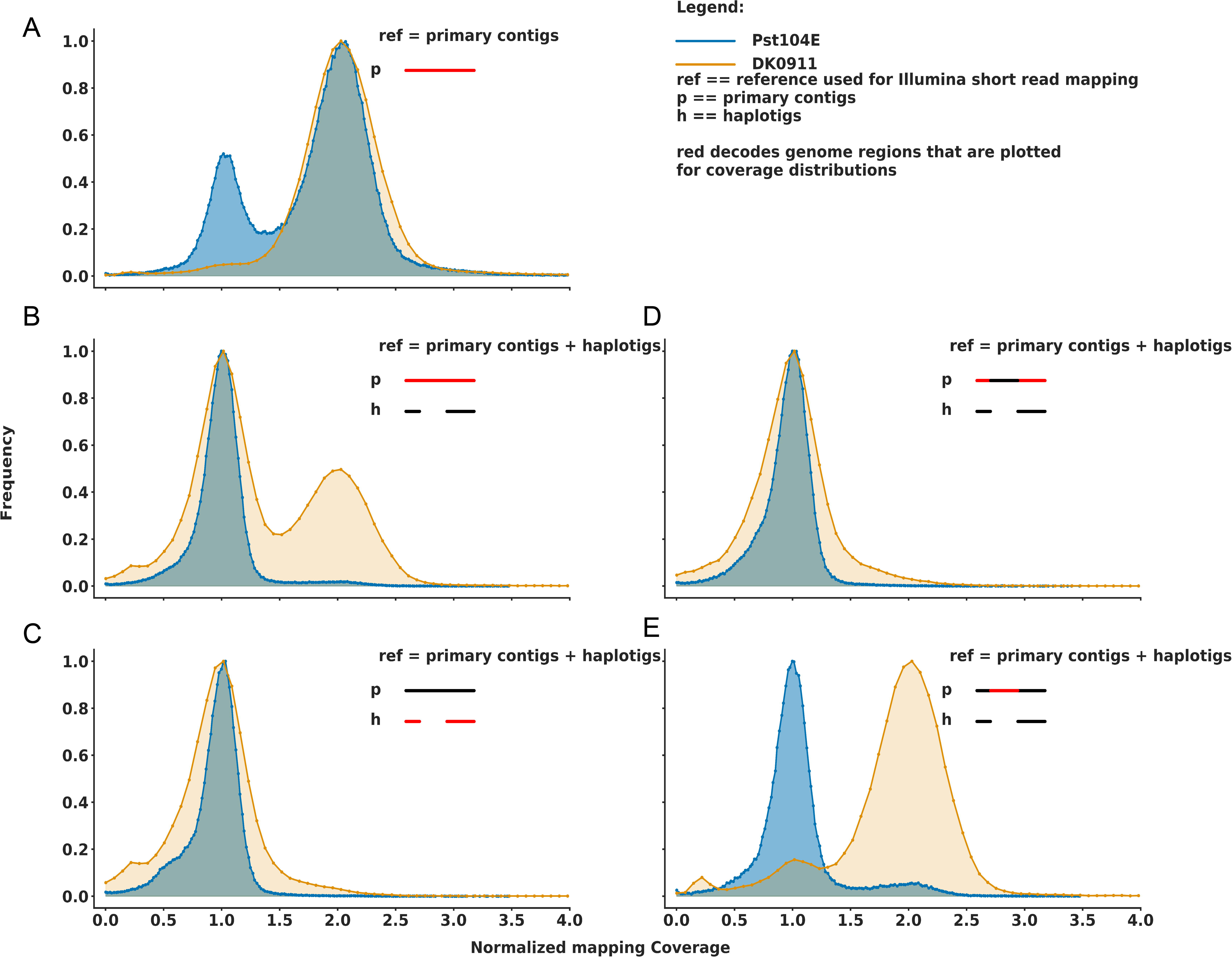
Low level of hemizygous regions in Pst-DK0911. Normalized Illumina short-read coverage plots for the red color-coded genome regions in primary contigs when mapping against primary contigs only (A) or when mapping against primary contigs and haplotigs (B, D, and E). (C) shows the coverage of haplotigs when mapping against primary contigs and haplotigs. Coverage was normalized to one in regions of the genome that are fully phased. The Pst-DK0911 contains significantly less hemizygous regions when compared to Pst-104E as it only has a single 2x coverage peak when mapping against primary contigs only (A). In addition, nearly all regions in primary contigs that do not possess a corresponding haplotig have 2x coverage suggesting that these regions are collapsed homozygous regions in the assembly and not hemizygous (E).

**Figure 3:**
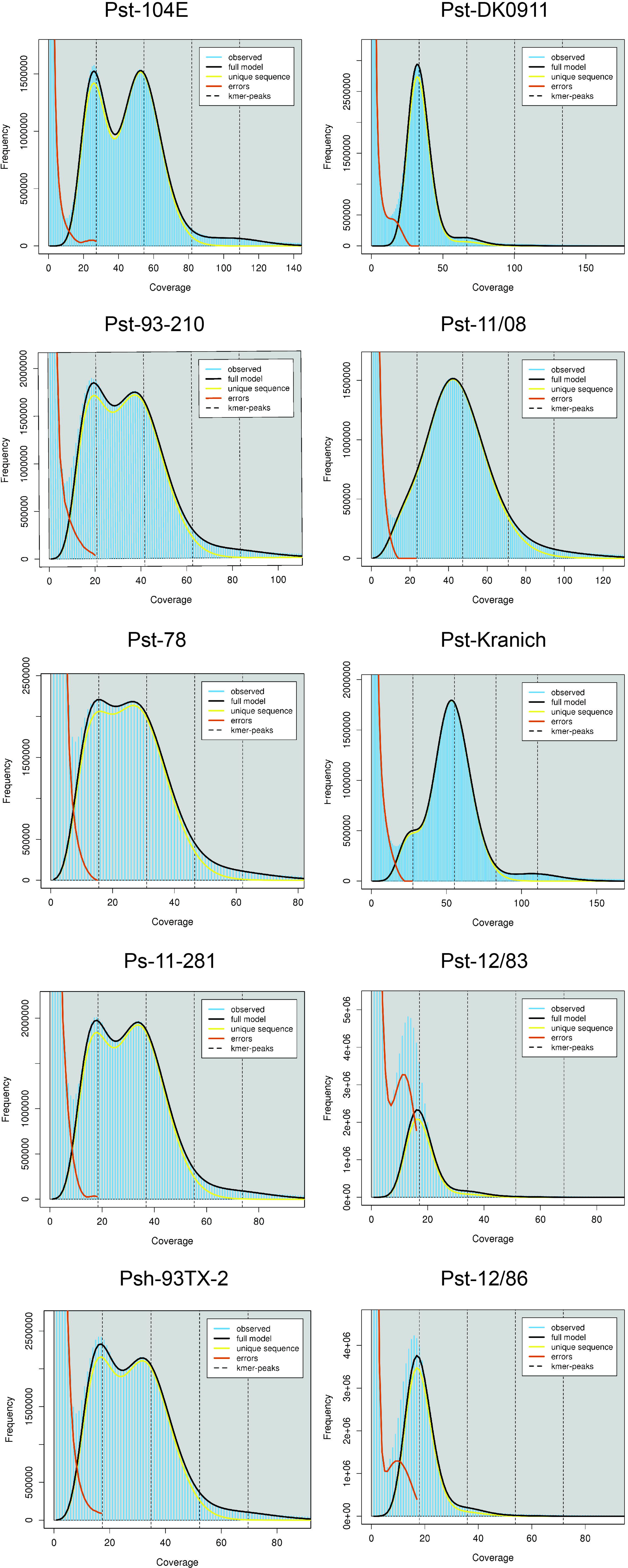
Low levels of heterozygosity in four additional *P*. *striiformis* f. sp*. tritici* isolates linked to the Himalayan region. K-mer frequency plots from Illumina short-read raw data generated with jellyfish and GenomeScope2. High levesl of heterozygosity are exemplified by double peaks in the k-mer frequency profile as shown for isolates in the left column including Pst-104E, Pst-93-210, and Pst-78. High levels of homozygosity are exemplified by a single peak in the k-mer frequency profile as shown for isolates in the right column including Pst-DK0911, Pst-11/08, and Pst-Kranich. All isolates in the right column are directly linked to the recent incursion event derived from the Himalayan region. The x-axis displays k-mer coverage and is dependent on the sequencing depth of the Illumina sequencing library. See Supplemental Data 5 for SRA identifiers and links to full GenomeScope2 reports.

**Figure 4:**
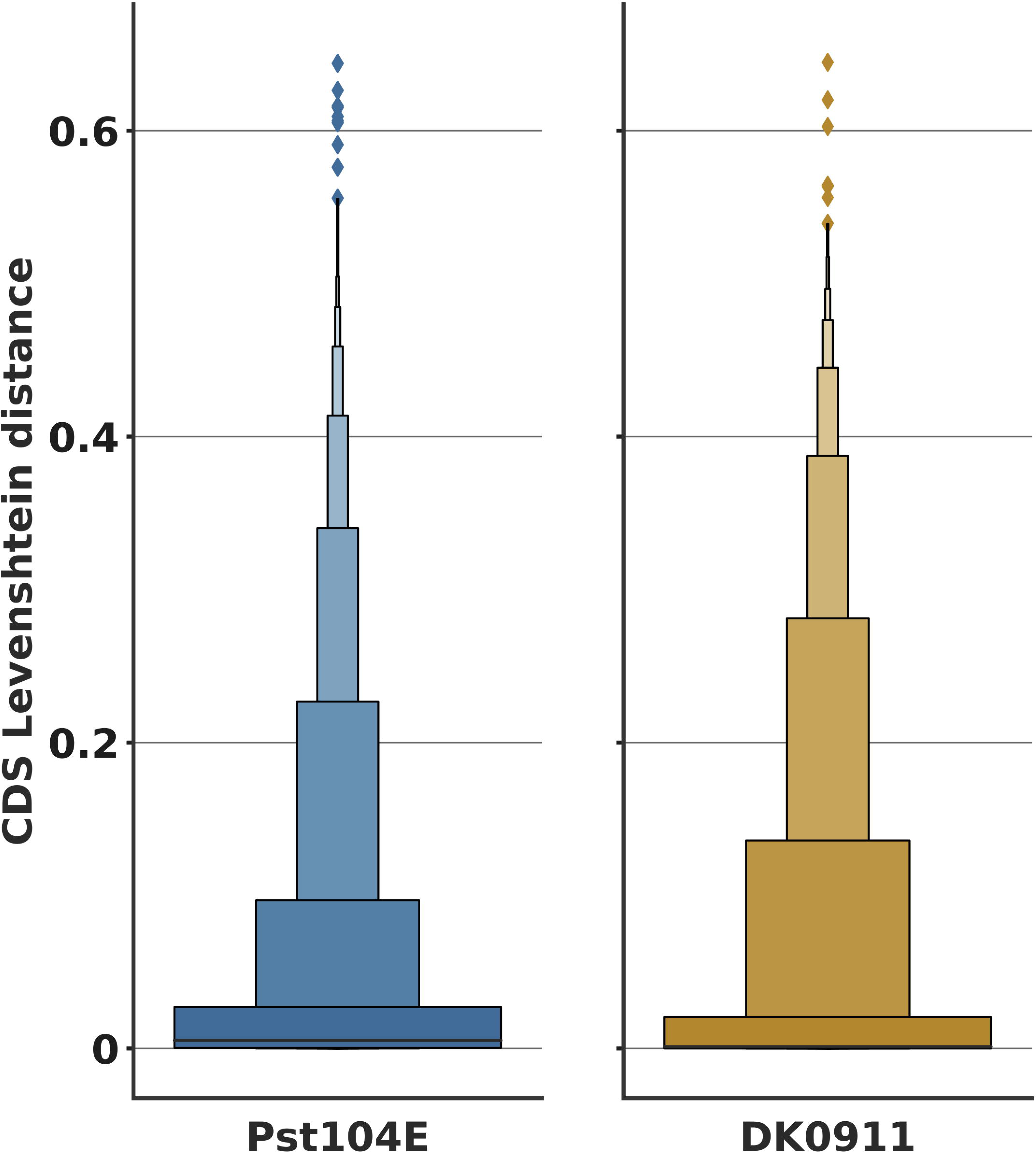
Different level of allelic gene pair variation in Pst-DK0911. Boxenplot of levenshstein edit distance of coding regions of high-quality allelic gene pairs in Pst-104E and Pst-DK0911. For values greater than zero, n[Pst-104E]=7472 and n[Pst-DK0911]=4344.

### Reduced telomere length in Pst-104E might be caused by long-term asexual reproduction

It is well known that telomere ends shorten with increasing mitotic cell divisions due to the incomplete replication of chromosome ends in mammalian cells (Aubert & Lansdorp 2008) but this phenomenon is less studied in filamentous fungi (Wang et al. 2014). We tested if telomere length is different in Pst-104E compared to Pst-DK0911 assuming a haploid chromosome number of 18 based on the observation that closely related rust fungi such as *P. graminis* f. sp. *tritici* (Boehm et al. 1992; Sperschneider et al. 2020; Li et al. 2019) and *Melampsora lini* (Boehm 1992) have 18 chromosomes in each nucleus. Our hypothesis was that if Pst-104E has been asexual for a longer time period and hence undergone a greater number of successive mitotic cell divisions, it should have shorter telomeres than Pst-DK0911. We approximated telomere length via searching the canonical ‘TTAGGG’ telomere repeat and variants thereof in whole genome Illumina short-read data (see Methods). We estimated 55 telomere repeats in Pst-DK0911 and 18 in Pst-104E, which suggests significant telomere shortening in Pst-104E. Consistently, the canonical ‘TTAGGG’ repeat accounts only for 76% of all patterns in Pst-104E versus 84% in Pst-DK0911 (see Supplemental Data 9 for full analysis results). This suggesting that Pst-104E has accumulated more variations at its telomeres over time. Overall, this observation of relatively short telomeres in Pst-104E compared to Pst-DK0911 is consistent with hypothesis of increased mitotic cell division as suggested by the increased heterozygosity in Pst-104E.

### Inter-lineage genome variation is dominated by variation in transposable elements

Next we tested for variation and presence-absence polymorphisms between the Pst-104E and Pst-DK0911 genomes using a conservative coverage based read mapping approach. We defined unique regions (highly variable or absent) in one genome if they fell below a 10% normalized coverage threshold in cross-mapping (e.g. Pst-104E reads onto the Pst-DK0911 genome), but not self-mapping (e.g. Pst-104E reads on to the Pst-104E genome) experiments. In total, we identified 6 Mb of sequence that was unique to Pst-DK0911 and 21 Mb to Pst-104E. We then considered how many genes are completely contained in these unique regions. We identified 724 variable genes that were in the Pst-DK0911 genome and not covered by Pst-104E reads and hence absent in Pst-104E. Conversely, we identified 2394 variable genes of Pst-104E that were not covered by Pst-DK0911 reads and hence missing from Pst-DK0911. When analyzing if any specific gene group is enriched or depleted within this variable gene set, we found that BUSCOs are significantly depleted in both cases (Multiple testing corrected Fisher’s exact test p-value = 3.1 e-11 for Pst-DK0911 and p-value = 8.2 e-9 for Pst-104E as reference), which is expected for genes that belong to the highly conserved core genome. We could not find any significant enrichment signal for candidate effectors or secreted proteins in either Pst-104E or Pst-DK0911. On the contrary, genes encoding for secreted proteins appeared to be depleted from the Pst-104E variable gene set (multiple testing corrected Fisher’s exact test p-value = 2.2 e-5).

We next tested if the unique regions were randomly distributed across the genomes or if they contained a different number of genes than expected by chance. To do this, we performed a permutation test by randomly reshuffling the unique regions to establish the expected distribution of variable genes when the unique regions are randomly distributed. In both cases BUSCO genes were significantly depleted from unique regions (Supplemental Figure 7). In the case of Pst-DK0911, candidate effectors and genes encoding secreted proteins were also depleted from unique regions. In contrast, more genes than expected by chance were fully contained within the unique genome regions of Pst-DK0911 and hence absent from Pst-104E. The reverse analyses did not reveal any strong signal for any gene group except for genes encoding for BUSCOs and secreted proteins which were significantly depleted from unique regions in Pst-104E. Next we tested if TEs contribute to sequence variation between the two genomes using a similar approach (see methods for details). Indeed, TEs were significantly enriched in unique regions identified by our coverage based read mapping approach (Supplemental Figure 8 A and B). As read mapping is known to suffer in repetitive regions, we performed identical tests based on whole genome alignments allowing for multi-mapping with a percentage identity coverage cut-off of 95%. Consistently, this second independent method also identified that TEs are highly enriched in unique regions in both genomes (Supplemental Figure 9 A and B). Overall, these analyses suggest that sequence variation between the two isolates is mostly caused by variation in TE sequences.

### Inter-lineage gene content variation does not favor candidate effectors

Similar to previous observations in *P*. *striiformis* f. sp*. tritici* (Schwessinger et al. 2018; Xia et al. 2018) and other rust fungi (Duplessis et al. 2011; Miller et al. 2018), candidate effectors did not show any obvious genome compartmentalization in Pst-DK0911 (Supplemental Figure 10). Candidate effectors were not located in gene sparse regions or linked to TEs. Yet similar to Pst-104E, candidate effectors appeared to be slightly closer to BUSCOs than to other genes (Supplemental Figure 10). Next, we tested if candidate effectors are more variable between the two isolates compared to all genes. We performed gene family (orthology) analysis with OrthoFinder (Emms & Kelly 2018) using 59 fungal proteomes representing 38 fungal species with a focus on rust fungi (see Supplemental Data 6 for details). Overall, candidate effectors did not appear to be significantly enriched in the singleton gene pool of either isolate. When identifying genes without orthologs in one of the two isolates, a total of 1275 and 1404 proteins were singletons in Pst-DK0911 and in Pst-104E, respectively. As expected BUSCOs are depleted from the singleton sets in both genomes (multiple testing error corrected Fisher’s exact test p-value = 3.0 e-33 for Pst-DK0911 and p-value = 5.3 e-31 for Pst-104E). When we investigated the EffectorP predicted candidate effector subset in Pst-104E we saw a weak signal of enrichment in the singleton gene set (Multiple testing error corrected Fisher’s exact test p-value = 1.0 e-4). In contrast, genes belonging to the secretome were depleted from the singleton gene set in Pst-DK0911 (multiple testing error corrected Fisher’s exact test p-value = 4.6 e-07).

### Candidate effectors fall into conserved and hypervariable categories

While candidate effectors were not specifically enriched in presence-absence polymorphisms between Pst-DK0911 and Pst-104E, we tested if some of the candidate effectors displayed signs of positive selection. First, we investigated the variation between syntenic orthologs on the nucleotide and amino acid level. About 44% and 57% of all effector candidates displayed variation at the nucleotide level, which is slightly below the genome average of 65% in Pst-DK0911 and 70% in Pst-104E, respectively. The variation between candidate effector genes was higher than BUSCOs but similar to the overall gene pool (Figure 5 A, see Supplemental Data 10-13 for statistical testing and gene category annotation). This trend was very similar at the protein level with 36% and 52% of candidate effectors displaying variation in Pst-DK0911 and Pst-104E, respectively. The variation in BUSCOs was significantly reduced compared to all other gene groups, yet similar between all other gene groups. This suggests that candidate effectors are not distinct from other gene pairs showing overall similar levels in sequence conservation between isolates. Lastly, we calculated the dN:dS ratio for all syntenic ortholog gene pairs. Indeed, 51 and 133 candidate effectors displayed a dN:dS ratio greater than one and hence a strong signal for positive selection in Pst-DK0911 and Pst-104E, respectively.

**Figure 5:**
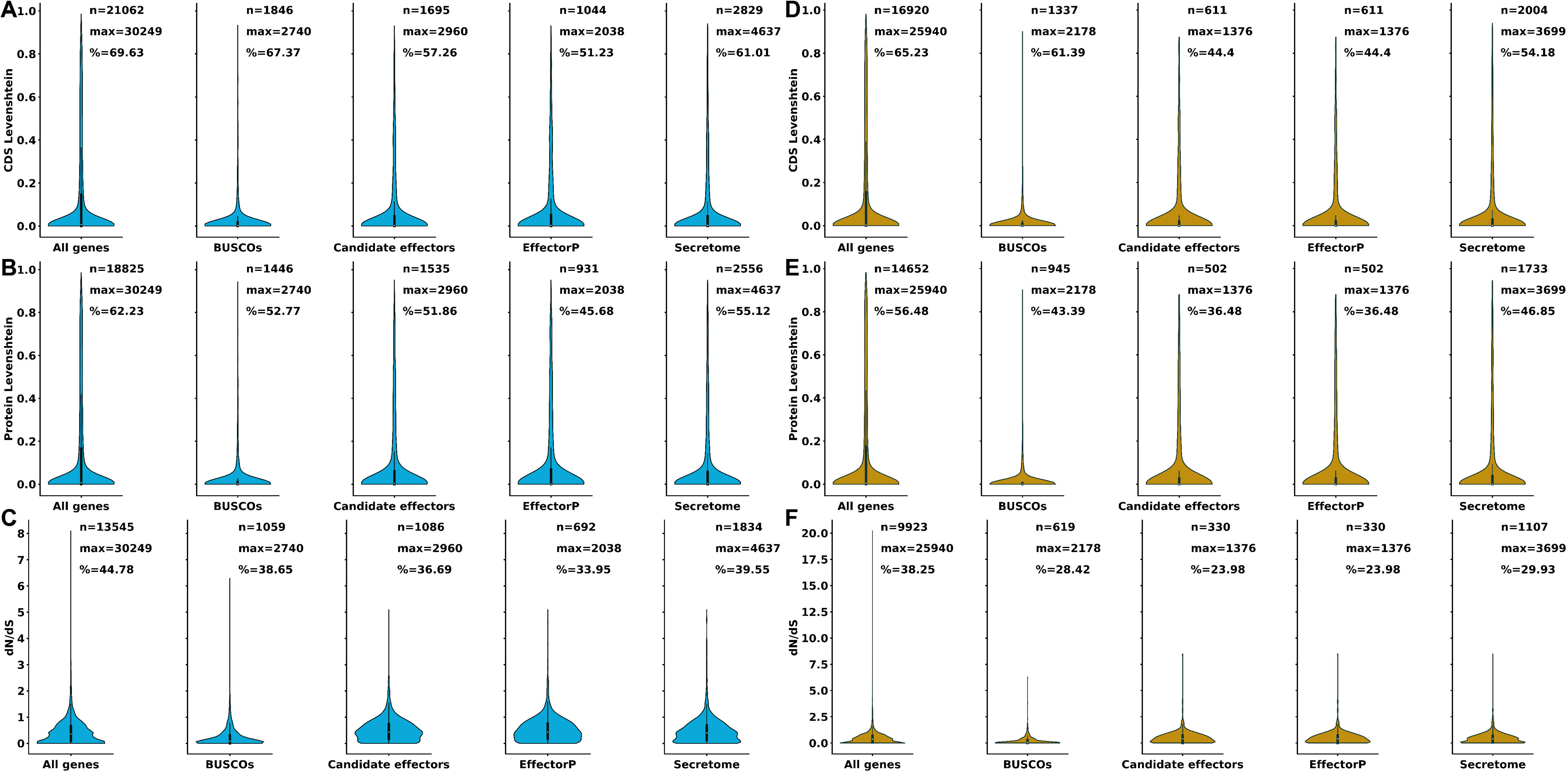
A subset of effector candidates displays patterns of positive selection. Violinplots of the pairwise comparison of syntenic orthologs in Pst-104E and Pst-DK0911 split according to gene classifications. (A) Shows the Levenshtein distance at the coding sequence level (CDS). (B) Shows the Levenshtein distance at the protein level. (C) Shows the dN:dS ratio. n indicates number of orthologs genes or proteins with any difference. max indicates the total number of orthologous genes or proteins % indicates the percentage of orthologs genes or proteins with any variation greater than zero. See the methods section for the definition of different gene categories. See Supplemental Data 10 and 11 for detailed statistical analysis of the complete data set.

### Candidate effectors do not show strong signals for altered synteny

While candidate effectors are not compartmentalized in rust fungi, we aimed to explore other signals of specific genome organization in regard to candidate effectors in *P*. *striiformis* f. sp*. tritici*. The availability of two highly contiguous partially phased genomes enabled us to ask the question if candidate effectors displayed an altered synteny pattern similar to *R* genes in plants. *R* genes in plants are the least syntenic genes in land plants (Zhao & Schranz 2019). If this were the case for effectors, it could indicate that candidate effectors are in genomic regions with higher recombination rates or general higher plasticity leading to disruption of gene synteny. Using a similar approach as described for syntenic gene orthologs in plants (Zhao & Schranz 2019), we first tested how sensitive different gene groups are towards allowing a varying number of gene gaps within synteny blocks using primary contigs as queries against the complete assembly as subject. As expected (Zhao & Schranz 2019), BUSCOs were the least affected by the number of gaps (Figure 6). The number of BUSCOs within a synteny block plateaued at about 97% already at a maximum gap size of two or three in Pst-DK0911 or Pst-104E, respectively (Figure 6 A). In contrast, for all other gene groups the percentage of genes contained in synteny blocks kept on increasing until the maximum tested allowed gap size of 35. Over 94% and 91% of all candidate effectors were contained within synteny blocks with a gap size of 8 or more in Pst-DK0911 or Pst-104E, respectively. This is more than the genome average of 88% and 86%, respectively. Hence overall candidate effectors do not appear to be less syntenic than the genome average. This observation was further supported by the fact that we observed very little difference in the mean best synteny block length at various numbers of allowed gaps within the alignments (Figure 6 B). While BUSCOs appeared to have slightly longer mean synteny block length, the variation of synteny block length was significant in each gene group (see Supplemental Figure 11) not allowing for any generalization. Overall, these observations are consistent with the fact that candidate effectors are also not enriched in inter-lineage singletons and appear to be mostly surprisingly conserved at the gene level.

**Figure 6:**
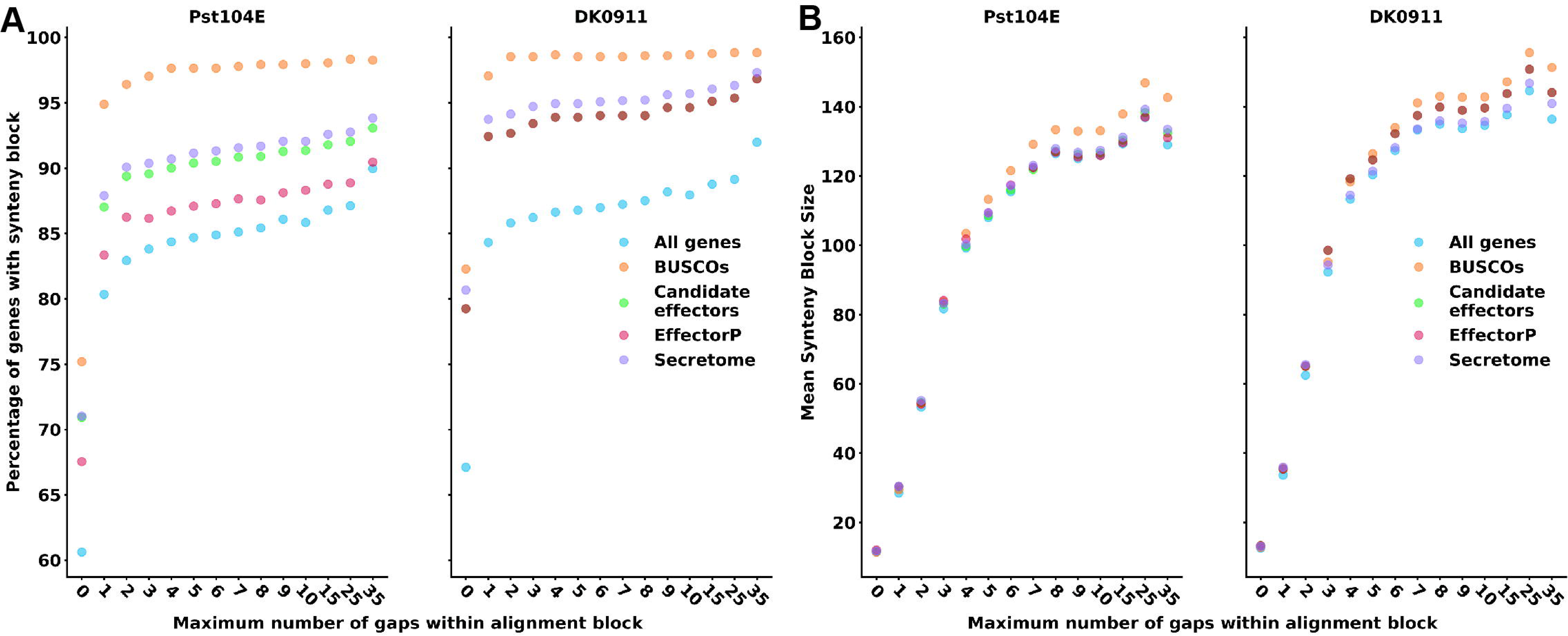
Candidate effectors are mostly syntenic between the two *P*. *striiformis* f. sp*. tritici* isolates. (A) Shows the percentage of genes that are syntenic at increasing number of allowed gaps within the synteny block. (B) Shows the mean synteny block length at increasing number of allowed gaps within the synteny block. The minimum synteny block size is three. Data points only include genes on the primary contig of the query genome. Data points represent different gene classifications. See the methods section for the definition of different gene categories. See Supplemental Figure 2 explaining the influence of gap size on synteny blocks.

## Discussion

### Genomic insights into the genetic effects of sexual reproduction in *P*. *striiformis* f. sp*. tritici*

In the early 2010s, *P*. *striiformis* f. sp*. tritici* isolates from the ‘Warrior’ (PstS7) lineage arrived in Europe severely impacting wheat production by causing stripe rust on previously resistant wheat varieties (Hovmøller et al. 2016; Hubbard et al. 2015). The Pst-DK0911 isolate analyzed in this manuscript belongs to the PstS7 lineage and was captured in Denmark in June 2011 (Hovmøller et al. 2016). The PstS7 lineage is genetically distinct from the pre-existent globally important lineages such as PstS0 and PstS1/2, which are part of presumed asexual lineages at least tracing back to isolates in Europe in the 1950s (Thach et al. 2016) and Africa in the 1980s (Walter et al. 2016), respectively. Pst-DK0911 in contrast is closely related to sexual *P*. *striiformis* f. sp*. tritici* populations in the Himalayan regions, which represent the center of genetic diversity for *P*. *striiformis* f. sp*. tritici* (Ali et al. 2017; Ali et al. 2014). Pst-DK0911 is one of very few reported *P*. *striiformis* f. sp*. tritici* isolates that has undergone the complete sexual cycle under laboratory conditions (Rodriguez-Algaba et al. 2014). Self-crosses of Pst-DK0911 suggest that this isolate is largely homozygous, in terms of having two recognized alleles, for several recognized effector loci as none of the tested progeny isolates were able to infect wheat differential lines that were resistant to the parental isolate. Hence, we hypothesize that Pst-DK0911 originated from a highly inbred population that potentially might have undergone several rounds of selfing or crosses between highly related genotypes leading to relatively low heterozygosity. Indeed, our data supports this hypothesis as Pst-DK0911 displays the lowest heterozygosity at the SNP level of any *P*. *striiformis* f. sp*. tritici* isolate analyzed so far and four additional isolates recently arrived from the Himalayan region show similar low heterozygosity levels based on k-mer frequency plots. In addition, Pst-DK0911 displays about half the inter-haplotype structural variation when compared to Pst-104E, which belongs to the long-term asexual lineage PstS0. This observation is consistent with the insight gained from *Puccinia coronata* f. sp. *avenae* where the genome of an isolate closely related to sexual populations displays reduced heterozygosity by comparison to the genome of an isolate more closely related to long-term asexual populations (Miller et al. 2018). It is likely that repeated sexual cycles of *P*. *striiformis* f. sp*. tritici* lead to reduced heterozygosity as the physiology of sexual reproduction favors inbreeding within one population. In addition, sexual recombination allows for gene conversion and unequal crossover at homologous sides including transposable elements (Underwood & Choi 2019; Liu et al. 2018). Interestingly, the variation analysis of high-quality gene-pairs in Pst-DK0911 revealed that its genome harbors discontinuous populations of gene-pairs. In general, variation in gene-pairs in dikaryotic rust fungi can be caused by two major factors. The first would be continuous accumulation of mutations during asexual reproduction cycles leading to linear accumulation of variation between gene-pairs. The second being variation that has been introduced at a given point in time, e.g. mating of two genetically distinct parents or somatic recombination via the transfer of single nucleus. The gene pair variation in Pst-DK0911 appears to be shaped by both sources of variation and is best explained by an initial cross of genetically distinct parents, several rounds of selfing of closely related individuals on barberry, followed by a limited number of asexual reproduction cycles on wheat.

### The high quality mitochondrial genome of *P*. *striiformis* f. sp*. tritici* DK0911 reveals long low-complexity AT regions

The mitochondrial genome of Pst-DK0911 consists of extensive intergenic regions interspersed with long low-complexity AT repeats. The overall structure and gene content of the Pst-DK0911 is consistent with a recently high-quality mitochondrial genome of Pst-CYR32 (Li et al. 2020). Comparative analysis of *P*. *striiformis* f. sp*. tritici* mt-genome with rust and other fungi revealed that these repeats are unique to *P*. *striiformis* f. sp*. tritici* mt-genome. The overall GC content of the *P*. *striiformis* f. sp*. tritici* mt-genome is 31% rising to 35.8% in the protein-coding regions. This is similar to other fungal mt-genomes (Torriani et al. 2014; Stone et al. 2010), although the *P*. *striiformis* f. sp*. tritici* mt-genome is comparatively large. The size of fungal mt-genomes can vary by as much as 10-fold; for example, the smallest reported fungal mt-genome is just 20,063 bp for *Candida glabrata* (Koszul et al. 2003) and the largest 230,000 bp for *Rhizoctonia solani* (Losada et al. 2014). The genetic content of these mt-genomes is largely conserved and consists of 14 genes with the order of the genes being highly variable between different fungi. This variability in gene order is the highest in basidiomycete fungi (Aguileta et al. 2014). The gene order of *P*. *striiformis* f. sp*. tritici* mt-genome is similar to the close relative basidiomycete *Phakopsora sp.*, but the order of tRNA genes varies (Stone et al. 2010). Overall, this high-quality reference genome will enable the analysis of mitochondrial inheritance in *P*. *striiformis* f. sp*. tritici* in future studies.

### The potential effects of long-term asexual reproduction on genome expansion, heterozygosity and fitness in *P*. *striiformis* f. sp*. tritici*

In contrast to Pst-DK0911 (Rodriguez-Algaba et al. 2014), isolates belonging to the long-term asexual lineages of PstS0 and PstS1/2 are often compromised in teliospore production on wheat when compared to isolates from regions with signs of sexual reproduction such as China, Nepal, and Pakistan (Ali et al. 2010). Teliospore production is the entry point into the sexual infection cycle on barberry. It was hypothesized that this reduction is caused by counterselection of genes contributing to sexual reproduction in these two lineages given their likely exclusive asexual reproduction over multiple decades (Rodriguez-Algaba et al. 2014). Interestingly, our read cross-mapping presence-absence analysis identified a non-random distribution of genes located in highly variable regions of Pst-DK0911 compared to Pst-104E. These and other highly variable genes might be involved in sexual reproduction in Pst-DK0911. In the future, it will be interesting to investigate expression patterns of these genes during teliospore production on wheat and during the sexual cycle of *P*. *striiformis* f. sp*. tritici* on barberry.

Overall, our comparative analysis suggests that long-term asexual reproduction in Pst-104E has led to a continuous accumulation of mutations, structural variations and a high level of heterozygosity between the two dikaryotic nuclei. The observed gene-pair variation pattern in Pst-104E is distinct compared to Pst-DK0911 and best explained by the continuous accumulation of mutations during asexual reproduction without other major events introducing genetic diversity. Of course, we cannot exclude a somatic hybridization event in Pst-104E’s distant past similar to what has been recently suggested for *P*. *striiformis* (Lei et al. 2016a) and clearly demonstrated at the whole genome level for the *Puccinia graminis* f.sp. *tritici* isolate Ug99 (Li et al. 2019). In either case, the extensive relative telomere shortening in Pst-104E does support the hypothesis of an extensive number of mitotic cell divisions without interruption by meiosis as telomere shortening is an indication of aging in other organisms (Aubert & Lansdorp 2008). Long-term asexual reproduction and independent accumulation of mutations in both nuclei might have also impacted Pst-104E’s overall fitness. This effect of accumulation of mutations in asexual ‘diploids’ are often referred as the Meselson effect (Weir et al. 2016) and might at least in part explain the rapid replacement of the PstS0 lineage by novel incursion e.g. in Australia in the early 2000s (Wellings 2007).

Another striking fact is the relative genome size increase in Pst-104E which is partially (∼54%) explained by the expansion of the absolute TE content in this isolate. The contribution of TE expansion to genome size increase is well documented for plants, animals, and fungi. The overall TE family identity profile (TE ‘age’) is not different between Pst-104E and Pst-DK0911 with a most recent peak centering around 80% sequence identity at the family level. This indicates that TEs expansion in Pst-104E and Pst-DK0911 has happened in recent history. This is distinct to other fungal pathogen belonging to the class of *Leotiomycete* where major TE bursts can be observed at various TE ‘ages’. For example, there has been a very recent TE burst in *Blumeria graminis* f. sp. *hordei* DH14 with TE family level sequence identify of greater than 95%. At other end of the spectrum *Rhynchosporium commune* and *Rynchosporium secalis* only display a relative old TE burst with average sequence identify at the TE family level of around 65% (Frantzeskakis et al. 2018).

In our case, most of the TE content difference between the two isolate coincides with the major TE burst at around 80% TE family identity. The difference in TE content could be either explained by selective purging of TEs via unequal crossovers during meiosis (Shirleen Roeder 1983) in Pst-DK0911 in its recent past during sexual reproduction cycles. This hypothesis is most parsimonious with the observed data. Alternatively, the extended asexual reproduction cycles in Pst-104E might have led to an increase in TE content in the absence of TE purging and continuous TE activity during infection. The latter has been recently shown for *Zymoseptoria tritici* in which specific TE families are expressed during the infection of wheat (Fouché et al. 2020). The upregulation of specific TE families during infection was isolate specific, which could explain the difference of relative abundance of some TE families over others. Indeed, TEs are the most variable genomic regions between Pst-DK0911 and Pst-104E as measured at the superfamily level. In the future, it will be important to explore the expression of TEs in several *P*. *striiformis* f. sp*. tritici* isolates during the infection of wheat. In addition, it would be fascinating to perform whole genome comparative analysis of a time series of PstS0 isolates available in the Stubbs collection ranging from the 1950s to the 1990s (collected by R.W. Stubbs, Wageningen University) and later on expanded until today by the Global Rust Reference Center (Thach et al. 2015). These analyses could directly reveal the impact of long-term asexual reproduction on genome architecture, heterozygosity, and TE movement and plasticity.

### Effector evolution in *P*. *striiformis* f. sp*. tritici* and beyond

Clearly candidate effectors are not compartmentalized or linked to TEs in *P*. *striiformis* f. sp*. tritici* (Schwessinger et al. 2018; Xia et al. 2018). The availability of two haplotype-aware assemblies allowed us to test if candidate effectors behave differently compared to other genes in terms of synteny. Overall, candidate effectors did not behave strikingly differently from all genes. Yet comparison between candidate effectors and secreted proteins revealed that fewer candidate effectors, and especially EffectorP predicted candidates, are within a synteny block at any given allowed gap-size. This might suggest that these genes have a higher turn-over in regions of the genome that have increased recombination rates. Future studies capturing more of the genetic diversity of *P*. *striiformis* f. sp*. tritici* with haplotype aware assemblies will answer if this trend of reduced synteny of candidate effectors holds true in general. Overall, candidate effectors were well conserved between our two isolates and not enriched in presence absence polymorphisms. Yet we could observe two distinct populations of candidate effectors. About 70% of all candidate effectors were invariable between the two isolates. These might be core effectors required for plant infection of this obligate biotroph with a focus on wheat or not under selection by the plant immune system within the experienced environments. At the same time the remaining set of candidate effectors displayed an increased level of variation and dN:dS ratios, which is likely caused by specific selection pressures exerted by the plant immune system. This was especially visible in Pst-104E. This is consistent with the fact that Pst-104E is more heterozygous with more distinct candidate effector genes reducing the reciprocal mapping rate compared to Pst-DK0911. Overall the difference effector repertoire between Pst-104E and Pst-DK0911 might explain the differences in virulence profiles or aggressiveness observed between these two *P*. *striiformis* f. sp*. tritici* lineages (Ali et al. 2017).

### Evolution of wheat rust fungi

Genome compartmentalization of effectors is absent in rusts (Duplessis et al. 2011; Miller et al. 2018; Schwessinger et al. 2018; Xia et al. 2018), in most other obligate biotrophic plant pathogens including mildews of wheat and barley (Frantzeskakis et al. 2019; Müller et al. 2019; Fletcher et al. 2019; Frantzeskakis et al. 2018) and several other oomycete and fungal pathogens such as *Ramularia collo-cygni* (Stam et al. 2019; Frantzeskakis et al. 2019; Frantzeskakis et al. 2019). This suggests that the two-speed genome compartmentalization is not universal and less common than initially anticipated (Frantzeskakis et al. 2019). This poses the question of what other genetic mechanisms lead to variation in candidate effector complements within and between *P*. *striiformis* f. sp*. tritici* lineages. It has been shown recently that candidate effectors are enriched in recombination hotspots in *Zymoseptoria tritici* (Grandaubert et al. 2019) and *Blumeria graminis* f. sp. *tritici* (Müller et al. 2019). Similarly, recombination within sexual populations in the Himalayan region could lead to reshuffling of candidate effector loci in *P*. *striiformis* f. sp*. tritici.* This might bring together the most suitable candidate effectors for infection of wheat lines currently grown on large acreages. Indeed, we hypothesize that *P*. *striiformis* f. sp*. tritici* undergoes a two-step selection process. First, candidate effector allele combinations must be suited to the infection of modern wheat varieties in contrast to strains only adapted to local barberry populations and the proximal grasses, which are likely regular hosts of *P*. *striiformis* f. sp*. tritici* within the region. Second, once a *P*. *striiformis* f. sp*. tritici* lineage is adapted to modern wheat varieties it must overcome the specific resistance genes within currently grown wheat varieties (Ali et al. 2017). This second step of adaptation is driven by genetic variation introduced during the asexual reproduction cycle of wheat. The mutations in recognized effectors can be caused by SNPs, small insertions and deletions, larger structural variations and TE movement. Because it is presumed that both nuclei evolve independently during asexual reproduction, recognized effectors that are heterozygous to begin with are highly likely to be lost during this process. This effect is illustrated by the two cloned recognized effectors of *P. graminis* f. sp*. tritici*, AvrSr35 and AvrSr50, which were both heterozygous in the isolates they were identified in (Chen et al. 2017; Salcedo et al. 2017).

A third genetic mechanism of introducing variation is hybridization between distinct species or lineages (Stukenbrock 2016). Hybridization leads to novel allele combinations and previously heterozygous recessive alleles to be unearthed. Interspecies hybridization is important for the emergence of new plant pathogens as demonstrated for *Blumeria graminis* f. sp. *triticalae*, which arose from a cross between *B. graminis* f. sp. *tritici* and *B. graminis* f. sp. *secali* (Menardo et al. 2016). A special form of hybridization is the swapping of whole nuclei between different dikaryotic rust isolates (Park & Wellings 2012). This process is referred to as somatic hybridization and has long been speculated upon based on low resolution genetic and biochemical markers or novel virulence profiles emerging during co-infection of asexual hosts (Park & Wellings 2012; Lei et al. 2016b). Yet only very recently, somatic hybridization has been conclusively demonstrated to be important for wheat rust evolution under natural conditions. A recent study used completely haplotype phased assemblies to show that the hypervirulent *P. graminis* f. sp. *tritici* Ug99 isolate is a product of nuclei swapping between isolates of an old South African and a poorly described Iranian *P. graminis* f. sp. *tritici* lineage (Li et al. 2019). It is likely that similar somatic hybridization events occur in *P*. *striiformis* f. sp*. tritici.* Somatic hybridization of *P*. *striiformis* f. sp*. tritici* circumvents the first adaptation process on wheat as each haploid nucleus is likely already adapted to wheat. In addition, it can rapidly generate novel effector allele combinations and drive rapid *R* gene adaptation by introducing heterozygosity to the effector gene profile. In addition, the impact of somatic hybridization on generating hypervirulent wheat rust strains increases with the number of genetically distinct lineages within a given location. This furthers the biosecurity risk of higher rates of novel pathotype incursions in wheat growing areas globally.

## Conclusions

The future of wheat rust and *P*. *striiformis* genomics lies in generating complete haplotype phased genomes that assort each set of chromosomes into specific karyotypes. This will enable us to predict the potential of generating novel allele combinations during the asexual reproduction cycle via mutations and somatic hybridization. In addition, haplotype phased genomes will facilitate the cloning of recognized effectors. In combination, this will enable us to better detect specific recognized effector allele pair combinations using modern long-read DNA sequence approaches and enable us to better model the durability of certain *R* gene stacking approaches. Lastly, it will be important to survey *P*. *striiformis* populations in the Himalayan regions, which are not adapted to modern wheat varieties but to locally growing wheat and grasses. This will provide insight into the true genetic diversity of *P*. *striiformis* and enable us to catalog the entire genetic diversity of effectors. In addition, we might identify effectors that are important for the infection of wheat or those that are negatively selected for by modern wheat varieties.

## Acknowledgments

We thank Ben Auxier, Dr. Jeff Ellis, Dr. Remco Stam, and two anonymous reviewers for excellent comments and suggestions. We thank Dr. Diane Saunders for immediately sharing Illumina raw read data for Pst-12/83 and Pst-12/83. The research on Pst-DK0911 conducted at Aarhus University was funded by the Danish Innovation Fund (grant no. 11-116241, RUSTFIGHT). This work was supported by an Australian Research Council DECRA (DE150101897) and Future Fellowship (FT180100024) to BS.

## Data availability

All raw sequencing data and the assembled genomes can be found on NCBI under the BioSample ID SAMN12098061, BioProject ID PRJNA588102, the primary contigs with ID WXWY00000000, haplotigs with ID WXWX00000000, and mitochondrial genome with ID MN746374. Additional, annotation files and axillary genome files can be found on zenodo 10.5281/zenodo.3556840. Supplementary data files can be found on zenodo 10.5281/zenodo.3663113. Supplementary figures can be found on zenodo 10.5281/zenodo.3663111. All code related to this project can be found on github https://github.com/Team-Schwessinger/DK_0911-1 and has a zenodo doi of 10.5281/zenodo.3663116.

## Authors contribution

The experimental work on Pst-DK0911 was made by Y-J.C. and C.K.S. under the supervision of A.F.J and M.S.H. Preliminary bioinformatic analyses of short read Illumina genomic and transcriptomic data for Pst-DK0911 was made by Y-J.C and J.K.V. Provision of a genetically pure stock of Pst-DK0911 and virulence phenotyping of Pst-DK0911 was made by M.S.H. B.S. conducted all bioinformatic analysis except for the mitochondrial genome assembly. R. T. contributed to bioinformatic analysis. R. N. analyzed the mitochondrial genome. All authors contributed intellectually to this study. B.S wrote the manuscript with contributions from Y-J.C, A.F.J, R. N. and J. P. R. All authors commented on drafts of this manuscript and approved the final submission.

---

**Supplemental Figure 1: Specific TE expansion in Pst-104E are absent from Pst-DK0911**

(A) Mean identity distribution relative to the consensus sequence of TE families as a proxy of TE age in Pst-104E and Pst-DK0911. (B) Relative difference in TE coverage accumulation (0-1) in Pst-104E compared to Pst-DK0911 versus TE identity distribution as proxy of TE age. Genome coverage per TE superfamily elements in Pst-DK0911 (C) and Pst-104E (D) with a minimum mean identity of 80% and more than 50 copies. These TEs are most likely to contribute to current genome evolution. Bars are color-coded according to the order level.

**Supplemental Figure 2: Impact of gap number on synteny block length**

Blue and brown indicate different genomes. Triangles depict genes. Red lines indicate gene pairing between the genomes.

(A) The synteny block length of ‘Gene 1’ is five. This is independent of any allowed minimal gap number.

(B) The synteny block length of ‘Gene 1’ is four when allowing for no gaps or a gap number of one. The synteny block length of ‘Gene 1’ is five when allowing for a gap number of two and above.

**Supplemental Figure 3: Map of the Pst-DK0911 mt-genome**

The outer ring represents genes (coding and non-coding) and their respective positions in the genome. The yellow colour represents genes of nicotinamide adenine dinucleotide ubiquinone oxidoreductase complex, the pink colour represents genes of cytochrome c oxidase complex, the light green colour represents cytochrome b gene, dark green represents ATP synthase genes, brown represents ribosomal protein gene, navy blue represents tRNAs genes and red represents rRNAs genes. The inner ring representing GC content. The gaps in the inner ring representing AT abundant repeat.

**Supplemental Figure 4: Four high-quality *P*. *striiformis* f. sp*. tritici* mitochondrial genomes are overall highly conserved**

LASTZ alignment of the Pst-DK0911 genome sequence with three additional mitochondrial genome sequences from Pst-93-210 (CM009485), Psh-93TX-2 (CM009486), and Pst-CY32 (MH891489). Alignments were generated in Geneious.

**Supplemental Figure 5: *P*. *striiformis* f. sp*. tritici* characteristic transposable element distribution in Pst-DK0911**

Repetitive element annotation on primary contigs (A) and on haplotigs (B). Top panels show percentages of genome coverage for all repetitive elements and different subcategories. These include TEs of class I (RNA retrotransposons) and class II (DNA transposons), simple sequence repeats (SSR), and unclassifiable repeats (no Cat). Middle and bottom panels show percentages of genome coverage of class I and class II TEs categorized to class, order, and superfamily levels wherever possible. Repetitive elements were identified using the REPET pipeline, and classifications were inferred from the closest BLAST hit (see Materials and Methods). The TE order is color coded in each ClassI and ClassII TE plot.

**Supplemental Figure 6: Hemizygous regions are common in older western *P*. *striiformis* isolates**

Normalized Illumina short-read coverage plots when mapping against long-read primary genome assemblies of Pst-104E, Pst-DK0911, Pst-93-210 (SAMN08200485), Psh-93TX-2 (SAMN08200486), and Ps-11-281 (SBIN00000000). All genome assemblies except for Pst-DK0911 have significant number of hemizygous regions.

**Supplemental Figure 7: Short read cross-mapping based presence-absence polymorphisms between Pst-104E and Pst-DK0911 is not specifically enriched for candidate effectors**

Expected (violinplots) and observed (red line) number of gene completely contained in low coverage regions in cross-mapping experiments for Pst-DK0911 (A) or Pst-104E (B) as reference. Expected distributions were generated by randomly reshuffling the observed low coverage regions 5000 times and measuring the number of fully contained genes in the random low coverage regions. P-values are two-sided and calculated by comparing the observed number of contained genes to the expected distribution (see methods for details).

**Supplemental Figure 8: Short read cross-mapping based variation and presence-absence polymorphisms between Pst-DK0911 and Pst-104E is enriched for TEs**

Expected (violinplots) and observed (red line) number of bases pair regions of TEs contained in low coverage regions in cross-mapping experiments for Pst-DK0911 (A) or Pst-104E (B) as reference. Expected distributions were generated by randomly reshuffling the observed low coverage regions 2500 times and measuring the number of fully contained genes in the random low coverage regions. P-values are two-sided and calculated by comparing the observed number of contained genes to the expected distribution (see methods for details).

**Supplemental Figure 9: Whole genome alignment-based variation and presence-absence polymorphisms between Pst-DK0911 and Pst-104E is enriched for TEs**

Expected (violinplots) and observed (red line) number of bases pair regions of TEs contained in low coverage regions in whole genome alignment experiments for Pst-DK0911 (A) or Pst-104E (B) as reference. Expected distributions were generated by randomly reshuffling the observed low coverage regions 2500 times and measuring the number of fully contained genes in the random low coverage regions. P-values are two-sided and calculated by comparing the observed number of contained genes to the expected distribution (see methods for details).

**Supplemental Figure 10: Candidate effector are not found in gene spares regions and are not linked to TEs in Pst-DK0911**

(A) Nearest-neighbor gene distance density hexplots for three gene categories, including allgenes, BUSCOs, and candidate effectors. Each subplot represents a distance density hexplot with the log10 3’-flanking and 5’-flanking distance to the nearest-neighboring gene plotted along the x axis and y axis, respectively. (B) Violin plots for the log10 distance to the most proximal transposable element for genes in each category without allowing for overlap. (C) Violin plots for the log10 distance to the most proximal gene in the same category for subsamples of each category equivalent to the smallest category size (n = 814). (D) Violin plots for the minimum distance (log10) of candidate effectors and BUSCOs to each other or a random subset of genes (n = 814). The P values for panels B, C, and D were calculated using the Wilcoxon rank-sum test after correction for multiple testing (Bonferroni; alpha = 0.05) on the linear distance in bases.

**Supplemental Figure 11: Candidate effectors synteny block size distribution reflects the genome average.**

Violinplots of the mean synteny block length in (A) Pst-104E and (B) Pst-DK0911 and split according to gene classifications. The maximum allowed gap number is eight. Minimum synteny block length is three.

n indicates numbers of orthologs within a synteny block.

max indicates the total number of orthologs.

% indicates the percentage of orthologs within a synteny block.

## References

Aguileta G et al. 2014. High Variability of Mitochondrial Gene Order among Fungi. Genome Biol Evol. 6:451–465. doi: 10.1093/gbe/evu028.

Ali S et al. 2016. CloNcaSe: Estimation of sex frequency and effective population size by clonemate re-sampling in partially clonal organisms. Mol Ecol Resour. n/a-n/a. doi: 10.1111/1755-0998.12511.

Ali S, Gladieux P, Rahman H, et al. 2014. Inferring the contribution of sexual reproduction, migration and off-season survival to the temporal maintenance of microbial populations: a case study on the wheat fungal pathogen Puccinia striiformis f.sp. tritici. Mol Ecol. 23:603–617. doi: 10.1111/mec.12629.

Ali S, Gladieux P, Leconte M, et al. 2014. Origin, Migration Routes and Worldwide Population Genetic Structure of the Wheat Yellow Rust Pathogen Puccinia striiformis f.sp. tritici. PLoS Pathog. 10:e1003903. doi: 10.1371/journal.ppat.1003903.

Ali S et al. 2017. Yellow Rust Epidemics Worldwide Were Caused by Pathogen Races from Divergent Genetic Lineages. Front. Plant Sci. 8. doi: 10.3389/fpls.2017.01057.

Ali S, Leconte M, Walker A-S, Enjalbert J, de Vallavieille-Pope C. 2010. Reduction in the sex ability of worldwide clonal populations of Puccinia striiformis f.sp. tritici. Fungal Genetics and Biology. 47:828–838. doi: 10.1016/j.fgb.2010.07.002.

Altschul SF, Gish W, Miller W, Myers EW, Lipman DJ. 1990. Basic local alignment search tool. J. Mol. Biol. 215:403–410. doi: 10.1016/S0022-2836(05)80360-2.

Anikster Y. 1986. Teliospore Germination in Some Rust Fungi. Phytopathology. 76:1026. doi: 10.1094/Phyto-76-1026.

Aubert G, Lansdorp PM. 2008. Telomeres and aging. Physiol. Rev. 88:557–579. doi: 10.1152/physrev.00026.2007.

Bairoch A, Apweiler R. 2000. The SWISS-PROT protein sequence database and its supplement TrEMBL in 2000. Nucleic Acids Res. 28:45–48.

Bao W, Kojima KK, Kohany O. 2015. Repbase Update, a database of repetitive elements in eukaryotic genomes. Mobile DNA. 6:11. doi: 10.1186/s13100-015-0041-9.

Beddow JM et al. 2015. Research investment implications of shifts in the global geography of wheat stripe rust. Nature Plants. 15132. doi: 10.1038/nplants.2015.132.

Bendtsen JD, Nielsen H, von Heijne G, Brunak S. 2004. Improved prediction of signal peptides: SignalP 3.0. J. Mol. Biol. 340:783–795. doi: 10.1016/j.jmb.2004.05.028.

Bernt M et al. 2013. MITOS: Improved de novo metazoan mitochondrial genome annotation. Molecular Phylogenetics and Evolution. 69:313–319. doi: 10.1016/j.ympev.2012.08.023.

Boehm EWA. 1992. An Ultrastructural Pachytene Karyotype for *Melampsora lini*. Phytopathology. 82:1212. doi: 10.1094/Phyto-82-1212.

Boehm EWA et al. 1992. An ultrastructural pachytene karyotype for Puccinia graminis f.sp. tritici. Can. J. Bot. 70:401–413. doi: 10.1139/b92-054.

Buchfink B, Xie C, Huson DH. 2015. Fast and sensitive protein alignment using DIAMOND. Nature Methods. 12:59–60. doi: 10.1038/nmeth.3176.

Bueno-Sancho V et al. 2017. Pathogenomic Analysis of Wheat Yellow Rust Lineages Detects Seasonal Variation and Host Specificity. Genome Biol Evol. 9:3282–3296. doi: 10.1093/gbe/evx241.

Cantu D et al. 2013. Genome analyses of the wheat yellow (stripe) rust pathogen Puccinia striiformis f. sp. tritici reveal polymorphic and haustorial expressed secreted proteins as candidate effectors. BMC Genomics. 14:270. doi: 10.1186/1471-2164-14-270.

Cantu D et al. 2011. Next generation sequencing provides rapid access to the genome of Puccinia striiformis f. sp. tritici, the causal agent of wheat stripe rust. PLoS ONE. 6:e24230. doi: 10.1371/journal.pone.0024230.

Chen J et al. 2017. Loss of AvrSr50 by somatic exchange in stem rust leads to virulence for Sr50 resistance in wheat. Science. 358:1607–1610. doi: 10.1126/science.aao4810.

Chen W, Wellings C, Chen X, Kang Z, Liu T. 2014. Wheat stripe (yellow) rust caused by Puccinia striiformis f. sp. tritici. Molecular Plant Pathology. 15:433–446. doi: 10.1111/mpp.12116.

Chin C-S et al. 2016. Phased diploid genome assembly with single-molecule real-time sequencing. Nat Meth. 13:1050–1054. doi: 10.1038/nmeth.4035.

Cuomo CA et al. 2016. Comparative Analysis Highlights Variable Genome Content of Wheat Rusts and Divergence of the Mating Loci. G3: Genes, Genomes, Genetics. g3.116.032797. doi: 10.1534/g3.116.032797.

Dale RK, Pedersen BS, Quinlan AR. 2011. Pybedtools: a flexible Python library for manipulating genomic datasets and annotations. Bioinformatics. 27:3423–3424. doi: 10.1093/bioinformatics/btr539.

Dobon A, Bunting DCE, Cabrera-Quio LE, Uauy C, Saunders DGO. 2016. The host-pathogen interaction between wheat and yellow rust induces temporally coordinated waves of gene expression. BMC Genomics. 17:380. doi: 10.1186/s12864-016-2684-4.

Duan X et al. 2010. Puccinia striiformis f.sp. tritici presents high diversity and recombination in the over-summering zone of Gansu, China. Mycologia. 102:44–53. doi: 10.3852/08-098.

Duplessis S et al. 2011. Obligate biotrophy features unraveled by the genomic analysis of rust fungi. PNAS. 108:9166–9171. doi: 10.1073/pnas.1019315108.

Edgar RC. 2004. MUSCLE: multiple sequence alignment with high accuracy and high throughput. Nucleic Acids Res. 32:1792–1797. doi: 10.1093/nar/gkh340.

Ellis JG, Lagudah ES, Spielmeyer W, Dodds PN. 2014. The past, present and future of breeding rust resistant wheat. Front. Plant Sci. 5:641. doi: 10.3389/fpls.2014.00641.

Emms DM, Kelly S. 2018. OrthoFinder2: fast and accurate phylogenomic orthology analysis from gene sequences. bioRxiv. 466201. doi: 10.1101/466201.

Fletcher K et al. 2019. Genomic signatures of heterokaryosis in the oomycete pathogen Bremia lactucae. Nature Communications. 10:2645. doi: 10.1038/s41467-019-10550-0.

Flutre T, Duprat E, Feuillet C, Quesneville H. 2011. Considering Transposable Element Diversification in De Novo Annotation Approaches. PLOS ONE. 6:e16526. doi: 10.1371/journal.pone.0016526.

Fouché S et al. 2020. Stress-Driven Transposable Element De-repression Dynamics and Virulence Evolution in a Fungal Pathogen. Mol Biol Evol. 37:221–239. doi: 10.1093/molbev/msz216.

Frantzeskakis L et al. 2018. Signatures of host specialization and a recent transposable element burst in the dynamic one-speed genome of the fungal barley powdery mildew pathogen. BMC Genomics. 19:381. doi: 10.1186/s12864-018-4750-6.

Frantzeskakis L, Németh MZ, et al. 2019. The Parauncinula polyspora Draft Genome Provides Insights into Patterns of Gene Erosion and Genome Expansion in Powdery Mildew Fungi. mBio. 10. doi: 10.1128/mBio.01692-19.

Frantzeskakis L, Kusch S, Panstruga R. 2019. The need for speed: compartmentalized genome evolution in filamentous phytopathogens. Molecular Plant Pathology. 20:3–7. doi: 10.1111/mpp.12738.

Garrison E, Marth G. 2012. Haplotype-based variant detection from short-read sequencing. arXiv:1207.3907 [q-bio]. http://arxiv.org/abs/1207.3907 (Accessed September 15, 2017).

Gladieux P et al. 2014. Fungal evolutionary genomics provides insight into the mechanisms of adaptive divergence in eukaryotes. Molecular Ecology. 23:753–773. doi: 10.1111/mec.12631.

Grabherr MG et al. 2011. Trinity: reconstructing a full-length transcriptome without a genome from RNA-Seq data. Nat Biotechnol. 29:644–652. doi: 10.1038/nbt.1883.

Grandaubert J, Dutheil JY, Stukenbrock EH. 2019. The genomic determinants of adaptive evolution in a fungal pathogen. Evolution Letters. 3:299–312. doi: 10.1002/evl3.117.

Grigoriev IV et al. 2014. MycoCosm portal: gearing up for 1000 fungal genomes. Nucleic Acids Res. 42:D699–704. doi: 10.1093/nar/gkt1183.

Haas BJ et al. 2008. Automated eukaryotic gene structure annotation using EVidenceModeler and the Program to Assemble Spliced Alignments. Genome Biology. 9:R7. doi: 10.1186/gb-2008-9-1-r7.

Harris RS. 2007. Improved Pairwise Alignment of Genomic Dna. PhD Thesis, Pennsylvania State University: USA.

Heitman J. 2015. Evolution of sexual reproduction: A view from the fungal kingdom supports an evolutionary epoch with sex before sexes. Fungal Biology Reviews. 29:108–117. doi: 10.1016/j.fbr.2015.08.002.

Hoff KJ, Lange S, Lomsadze A, Borodovsky M, Stanke M. 2016. BRAKER1: Unsupervised RNA-Seq-Based Genome Annotation with GeneMark-ET and AUGUSTUS. Bioinformatics. 32:767–769. doi: 10.1093/bioinformatics/btv661.

Hovmøller MS et al. 2016. Replacement of the European wheat yellow rust population by new races from the centre of diversity in the near-Himalayan region. Plant Pathol. 65:402–411. doi: 10.1111/ppa.12433.

Hovmøller MS, Justesen AF. 2007. Rates of evolution of avirulence phenotypes and DNA markers in a northwest European population of Puccinia striiformis f. sp. tritici. Molecular Ecology. 16:4637–4647. doi: 10.1111/j.1365-294X.2007.03513.x.

Hovmøller MS, Justesen AF, Brown JKM. 2002. Clonality and long-distance migration of Puccinia striiformis f.sp. tritici in north-west Europe. Plant Pathology. 51:24–32. doi: 10.1046/j.1365-3059.2002.00652.x.

Hovmøller MS, Yahyaoui AH, Milus EA, Justesen AF. 2008. Rapid global spread of two aggressive strains of a wheat rust fungus. Molecular Ecology. 17:3818–3826. doi: 10.1111/j.1365-294X.2008.03886.x.

Hubbard A et al. 2015. Field pathogenomics reveals the emergence of a diverse wheat yellow rust population. Genome Biology. 16:23. doi: 10.1186/s13059-015-0590-8.

Huerta-Cepas J et al. 2016. eggNOG 4.5: a hierarchical orthology framework with improved functional annotations for eukaryotic, prokaryotic and viral sequences. Nucleic Acids Res. 44:D286–D293. doi: 10.1093/nar/gkv1248.

Hunt M et al. 2015. Circlator: automated circularization of genome assemblies using long sequencing reads. Genome Biology. 16:294. doi: 10.1186/s13059-015-0849-0.

Jin Y, Szabo LJ, Carson M. 2010. Century-Old Mystery of Puccinia striiformis Life History Solved with the Identification of Berberis as an Alternate Host. Phytopathology. 100:432–435. doi: 10.1094/PHYTO-100-5-0432.

Jones P et al. 2014. InterProScan 5: genome-scale protein function classification. Bioinformatics. 30:1236–1240. doi: 10.1093/bioinformatics/btu031.

Kim D, Langmead B, Salzberg SL. 2015. HISAT: a fast spliced aligner with low memory requirements. Nat Meth. 12:357–360. doi: 10.1038/nmeth.3317.

Koren S et al. 2017. Canu: scalable and accurate long-read assembly via adaptive k-mer weighting and repeat separation. Genome Res. gr.215087.116. doi: 10.1101/gr.215087.116.

Korovesi AG, Ntertilis M, Kouvelis VN. 2018. Mt-rps3 is an ancient gene which provides insight into the evolution of fungal mitochondrial genomes. Molecular Phylogenetics and Evolution. 127:74–86. doi: 10.1016/j.ympev.2018.04.037.

Koszul R et al. 2003. The complete mitochondrial genome sequence of the pathogenic yeast Candida (Torulopsis) glabrata. FEBS Letters. 534:39–48. doi: 10.1016/S0014-5793(02)03749-3.

Krumsiek J, Arnold R, Rattei T. 2007. Gepard: a rapid and sensitive tool for creating dotplots on genome scale. Bioinformatics. 23:1026–1028. doi: 10.1093/bioinformatics/btm039.

Kurtz S et al. 2004. Versatile and open software for comparing large genomes. Genome Biol. 5:R12. doi: 10.1186/gb-2004-5-2-r12.

Lang BF, Laforest M-J, Burger G. 2007. Mitochondrial introns: a critical view. Trends in Genetics. 23:119–125. doi: 10.1016/j.tig.2007.01.006.

Laslett D, Canback B. 2004. ARAGORN, a program to detect tRNA genes and tmRNA genes in nucleotide sequences. Nucleic Acids Res. 32:11–16. doi: 10.1093/nar/gkh152.

Lechner M et al. 2011. Proteinortho: Detection of (Co-)orthologs in large-scale analysis. BMC Bioinformatics. 12:124. doi: 10.1186/1471-2105-12-124.

Lei Y et al. 2016a. Virulence and Molecular Characterization of Experimental Isolates of the Stripe Rust Pathogen (Puccinia striiformis) Indicate Somatic Recombination. Phytopathology^TM^. 107:329–344. doi: 10.1094/PHYTO-07-16-0261-R.

Lei Y et al. 2016b. Virulence and Molecular Characterization of Experimental Isolates of the Stripe Rust Pathogen (Puccinia striiformis) Indicate Somatic Recombination. Phytopathology. 107:329–344. doi: 10.1094/PHYTO-07-16-0261-R.

Li Chuang et al. 2020. The complete mitochondrial genomes of Puccinia striiformis f. sp. tritici and Puccinia recondita f. sp. tritici. Mitochondrial DNA Part B. 5:29–30. doi: 10.1080/23802359.2019.1674744.

Li Feng et al. 2019. Emergence of the Ug99 lineage of the wheat stem rust pathogen through somatic hybridisation. Nat Commun. 10:1–15. doi: 10.1038/s41467-019-12927-7.

Li H. 2013. Aligning sequence reads, clone sequences and assembly contigs with BWA-MEM. arXiv:1303.3997 [q-bio]. http://arxiv.org/abs/1303.3997 (Accessed September 14, 2017).

Li Yuxiang, Xia C, Wang M, Yin C, Chen X. 2019. Genome Sequence Resource of a Puccinia striiformis Isolate Infecting Wheatgrass. Phytopathology^TM^. 109:1509–1512. doi: 10.1094/PHYTO-02-19-0054-A.

Liu H et al. 2018. Tetrad analysis in plants and fungi finds large differences in gene conversion rates but no GC bias. Nat Ecol Evol. 2:164–173. doi: 10.1038/s41559-017-0372-7.

Lorrain C, Santos KCG dos, Germain H, Hecker A, Duplessis S. 2019. Advances in understanding obligate biotrophy in rust fungi. New Phytologist. 222:1190–1206. doi: 10.1111/nph.15641.

Losada L et al. 2014. Mobile elements and mitochondrial genome expansion in the soil fungus and potato pathogen Rhizoctonia solani AG-3. FEMS Microbiol Lett. 352:165–173. doi: 10.1111/1574-6968.12387.

Lowe TM, Eddy SR. 1997. tRNAscan-SE: A Program for Improved Detection of Transfer RNA Genes in Genomic Sequence. Nucleic Acids Res. 25:955–964. doi: 10.1093/nar/25.5.955.

Marçais G, Kingsford C. 2011. A fast, lock-free approach for efficient parallel counting of occurrences of k-mers. Bioinformatics. 27:764–770. doi: 10.1093/bioinformatics/btr011.

Markell SG, Milus EA. 2008. Emergence of a Novel Population of Puccinia striiformis f. sp. tritici in Eastern United States. Phytopathology. 98:632–639. doi: 10.1094/PHYTO-98-6-0632.

Maumus F, Quesneville H. 2014. Ancestral repeats have shaped epigenome and genome composition for millions of years in *Arabidopsis thaliana*. Nature Communications. 5:4104. doi: 10.1038/ncomms5104.

Mboup M et al. 2012. Genetic structure and local adaptation of European wheat yellow rust populations: the role of temperature-specific adaptation. Evolutionary Applications. 5:341–352. doi: 10.1111/j.1752-4571.2011.00228.x.

McKinney W. 2010. Data Structures for Statistical Computing in Python. In: pp. 51–56. http://conference.scipy.org/proceedings/scipy2010/mckinney.html (Accessed September 15, 2017).

Menardo F et al. 2016. Hybridization of powdery mildew strains gives rise to pathogens on novel agricultural crop species. Nat Genet. 48:201–205. doi: 10.1038/ng.3485.

Miller ME et al. 2018. De Novo Assembly and Phasing of Dikaryotic Genomes from Two Isolates of Puccinia coronata f. sp. avenae, the Causal Agent of Oat Crown Rust. mBio. 9:e01650–17. doi: 10.1128/mBio.01650-17.

Müller MC et al. 2019. A chromosome-scale genome assembly reveals a highly dynamic effector repertoire of wheat powdery mildew. New Phytologist. 221:2176–2189. doi: 10.1111/nph.15529.

Nattestad M, Schatz MC. 2016. Assemblytics: a web analytics tool for the detection of variants from an assembly. Bioinformatics. 32:3021–3023. doi: 10.1093/bioinformatics/btw369.

Nersisyan L, Arakelyan A. 2015. Computel: Computation of Mean Telomere Length from Whole-Genome Next-Generation Sequencing Data. PLoS One. 10. doi: 10.1371/journal.pone.0125201.

Park RF, Wellings CR. 2012. Somatic Hybridization in the Uredinales. Annual Review of Phytopathology. 50:219–239. doi: 10.1146/annurev-phyto-072910-095405.

Pertea M et al. 2015. StringTie enables improved reconstruction of a transcriptome from RNA-seq reads. Nat Biotech. 33:290–295. doi: 10.1038/nbt.3122.

Petersen TN, Brunak S, von Heijne G, Nielsen H. 2011. SignalP 4.0: discriminating signal peptides from transmembrane regions. Nat Meth. 8:785–786. doi: 10.1038/nmeth.1701.

Quesneville H et al. 2005. Combined Evidence Annotation of Transposable Elements in Genome Sequences. PLOS Computational Biology. 1:e22. doi: 10.1371/journal.pcbi.0010022.

Quinlan AR, Hall IM. 2010. BEDTools: a flexible suite of utilities for comparing genomic features. Bioinformatics. 26:841–842. doi: 10.1093/bioinformatics/btq033.

Radhakrishnan GV et al. 2019. MARPLE, a point-of-care, strain-level disease diagnostics and surveillance tool for complex fungal pathogens. BMC Biology. 17:65. doi: 10.1186/s12915-019-0684-y.

Ranallo-Benavidez TR, Jaron KS, Schatz MC. 2019. GenomeScope 2.0 and Smudgeplots: Reference-free profiling of polyploid genomes. bioRxiv. 747568. doi: 10.1101/747568.

Rapilly F. 1979. Yellow Rust Epidemiology. Annu. Rev. Phytopathol. 17:59–73. doi: 10.1146/annurev.py.17.090179.000423.

Rawlings ND, Barrett AJ, Finn R. 2016. Twenty years of the MEROPS database of proteolytic enzymes, their substrates and inhibitors. Nucleic Acids Res. 44:D343–D350. doi: 10.1093/nar/gkv1118.

Rodriguez-Algaba J, Walter S, Sørensen CK, Hovmøller MS, Justesen AF. 2014. Sexual structures and recombination of the wheat rust fungus Puccinia striiformis on Berberis vulgaris. Fungal Genetics and Biology. 70:77–85. doi: 10.1016/j.fgb.2014.07.005.

Salcedo A et al. 2017. Variation in the AvrSr35 gene determines Sr35 resistance against wheat stem rust race Ug99. Science. 358:1604–1606. doi: 10.1126/science.aao7294.

Savary S et al. 2019. The global burden of pathogens and pests on major food crops. Nature Ecology & Evolution. 1. doi: 10.1038/s41559-018-0793-y.

Schwessinger B et al. 2018. A Near-Complete Haplotype-Phased Genome of the Dikaryotic Wheat Stripe Rust Fungus Puccinia striiformis f. sp. tritici Reveals High Interhaplotype Diversity. mBio. 9:e02275–17. doi: 10.1128/mBio.02275-17.

Schwessinger B. 2017. Fundamental wheat stripe rust research in the 21(st) century. New Phytol. 213:1625–1631. doi: 10.1111/nph.14159.

Schwessinger B, Rathjen JP. 2017. Extraction of High Molecular Weight DNA from Fungal Rust Spores for Long Read Sequencing. In: Wheat Rust Diseases. Methods in Molecular Biology Humana Press, New York, NY pp. 49–57. doi: 10.1007/978-1-4939-7249-4_5.

Seudre O et al. 2018. Why outcross? The abandon-ship hypothesis in a facultative outcrossing/selfing fungal species. Fungal Genetics and Biology. 120:1–8. doi: 10.1016/j.fgb.2018.08.005.

Shirleen Roeder G. 1983. Unequal crossing-over between yeast transposable elements. Mol Gen Genet. 190:117–121. doi: 10.1007/BF00330332.

Simão FA, Waterhouse RM, Ioannidis P, Kriventseva EV, Zdobnov EM. 2015. BUSCO: assessing genome assembly and annotation completeness with single-copy orthologs. Bioinformatics. 31:3210–3212. doi: 10.1093/bioinformatics/btv351.

Singh RP et al. 2016. Disease Impact on Wheat Yield Potential and Prospects of Genetic Control. Annual Review of Phytopathology. 54:303–322. doi: 10.1146/annurev-phyto-080615-095835.

Sørensen CK, Hovmøller MS, Leconte M, Dedryver F, de Vallavieille-Pope C. 2014. New Races of Puccinia striiformis Found in Europe Reveal Race Specificity of Long-Term Effective Adult Plant Resistance in Wheat. Phytopathology. 104:1042–1051. doi: 10.1094/PHYTO-12-13-0337-R.

Sørensen CK, Thach T, Hovmøller MS. 2016. Evaluation of Spray and Point Inoculation Methods for the Phenotyping of Puccinia striiformis on Wheat. Plant Disease. PDIS-12-15-1477-RE. doi: 10.1094/PDIS-12-15-1477-RE.

Sperschneider J et al. 2015. EffectorP: predicting fungal effector proteins from secretomes using machine learning. New Phytol. n/a-n/a. doi: 10.1111/nph.13794.

Sperschneider J et al. 2020. The stem rust fungus Puccinia graminis f. sp. tritici induces waves of small RNAs with opposing profiles during wheat infection. bioRxiv. 469338. doi: 10.1101/469338.

Sperschneider J, Dodds PN, Gardiner DM, Singh KB, Taylor JM. 2018. Improved prediction of fungal effector proteins from secretomes with EffectorP 2.0. Mol. Plant Pathol. 19:2094–2110. doi: 10.1111/mpp.12682.

Stam R et al. A new reference genome shows the one-speed genome structure of the barley pathogen Ramularia collo-cygni. Genome Biol Evol. doi: 10.1093/gbe/evy240.

Steele KA, Humphreys E, Wellings CR, Dickinson MJ. 2001. Support for a stepwise mutation model for pathogen evolution in Australasian Puccinia striiformis f.sp. tritici by use of molecular markers. Plant Pathology. 50:174–180. doi: 10.1046/j.1365-3059.2001.00558.x.

Stone CL, Buitrago MLP, Boore JL, Frederick RD. 2010. Analysis of the complete mitochondrial genome sequences of the soybean rust pathogens Phakopsora pachyrhizi and P. meibomiae. Mycologia. 102:887–897. doi: 10.3852/09-198.

Stukenbrock EH. 2016. The Role of Hybridization in the Evolution and Emergence of New Fungal Plant Pathogens. Phytopathology. PHYTO-08-15-0184-RVW. doi: 10.1094/PHYTO-08-15-0184-RVW.

Suyama M, Torrents D, Bork P. 2006. PAL2NAL: robust conversion of protein sequence alignments into the corresponding codon alignments. Nucleic Acids Res. 34:W609–W612. doi: 10.1093/nar/gkl315.

Taylor JW, Hann-Soden C, Branco S, Sylvain I, Ellison CE. 2015. Clonal reproduction in fungi. PNAS. 112:8901–8908. doi: 10.1073/pnas.1503159112.

Testa AC, Hane JK, Ellwood SR, Oliver RP. 2015. CodingQuarry: highly accurate hidden Markov model gene prediction in fungal genomes using RNA-seq transcripts. BMC Genomics. 16:170. doi: 10.1186/s12864-015-1344-4.

Thach T, Ali S, Justesen A f., Rodriguez-Algaba J, Hovmøller M s. 2015. Recovery and virulence phenotyping of the historic ‘Stubbs collection’ of the yellow rust fungus Puccinia striiformis from wheat. Ann Appl Biol. 167:314–326. doi: 10.1111/aab.12227.

Thach T, Ali S, de Vallavieille-Pope C, Justesen AF, Hovmøller MS. 2016. Worldwide population structure of the wheat rust fungus Puccinia striiformis in the past. Fungal Genetics and Biology. 87:1–8. doi: 10.1016/j.fgb.2015.12.014.

Tillich M et al. 2017. GeSeq – versatile and accurate annotation of organelle genomes. Nucleic Acids Res. 45:W6–W11. doi: 10.1093/nar/gkx391.

Torriani SFF et al. 2014. Comparative analysis of mitochondrial genomes from closely related Rhynchosporium species reveals extensive intron invasion. Fungal Genetics and Biology. 62:34–42. doi: 10.1016/j.fgb.2013.11.001.

Underwood CJ, Choi K. 2019. Heterogeneous transposable elements as silencers, enhancers and targets of meiotic recombination. Chromosoma. 128:279–296. doi: 10.1007/s00412-019-00718-4.

Walker BJ et al. 2014. Pilon: An Integrated Tool for Comprehensive Microbial Variant Detection and Genome Assembly Improvement. PLOS ONE. 9:e112963. doi: 10.1371/journal.pone.0112963.

Walter S et al. 2016. Molecular markers for tracking the origin and worldwide distribution of invasive strains of Puccinia striiformis. Ecol Evol. n/a-n/a. doi: 10.1002/ece3.2069.

Wang N et al. 2014. Novel Telomere-Anchored PCR Approach for Studying Sexual Stage Telomeres in Aspergillus nidulans. PLOS ONE. 9:e99491. doi: 10.1371/journal.pone.0099491.

Wang Y et al. 2012. MCScanX: a toolkit for detection and evolutionary analysis of gene synteny and collinearity. Nucleic Acids Res. 40:e49–e49. doi: 10.1093/nar/gkr1293.

Weir W et al. 2016. Population genomics reveals the origin and asexual evolution of human infective trypanosomes. eLife Sciences. 5:e11473. doi: 10.7554/eLife.11473.

Wellings CR. 2011. Global status of stripe rust: a review of historical and current threats. Euphytica. 179:129–141. doi: 10.1007/s10681-011-0360-y.

Wellings CR. 2007. Puccinia striiformis in Australia: a review of the incursion, evolution, and adaptation of stripe rust in the period 1979–2006. Aust. J. Agric. Res. 58:567–575.

Wellings CR, McINTOSH RA. 1990. Puccinia striiformis f.sp. tritici in Australasia: pathogenic changes during the first 10 years. Plant Pathology. 39:316–325. doi: 10.1111/j.1365-3059.1990.tb02509.x.

Xia C, Wang M, Yin C, Cornejo OE, Hulbert Scot, et al. 2018. Genome sequence resources for the wheat stripe rust pathogen (Puccinia striiformis f. sp. tritici) and the barley stripe rust pathogen (Puccinia striiformis f. sp. hordei). MPMI. doi: 10.1094/MPMI-04-18-0107-A.

Xia C, Wang M, Yin C, Cornejo OE, Hulbert Scot H., et al. 2018. Genomic insights into host adaptation between the wheat stripe rust pathogen (Puccinia striiformis f. sp. tritici) and the barley stripe rust pathogen (Puccinia striiformis f. sp. hordei). BMC Genomics. 19:664. doi: 10.1186/s12864-018-5041-y.

Yang Z, Nielsen R. 2000. Estimating synonymous and nonsynonymous substitution rates under realistic evolutionary models. Mol. Biol. Evol. 17:32–43.

Yin Y et al. 2012. dbCAN: a web resource for automated carbohydrate-active enzyme annotation. Nucleic Acids Res. 40:W445–451. doi: 10.1093/nar/gks479.

Zeyl C. 2009. The role of sex in fungal evolution. Current Opinion in Microbiology. 12:592–598. doi: 10.1016/j.mib.2009.09.011.

Zhao T, Schranz ME. 2019. Network-based microsynteny analysis identifies major differences and genomic outliers in mammalian and angiosperm genomes. PNAS. 201801757. doi: 10.1073/pnas.1801757116.

Zheng W et al. 2013. High genome heterozygosity and endemic genetic recombination in the wheat stripe rust fungus. Nat Commun. 4:2673. doi: 10.1038/ncomms3673.

2019. PacificBiosciences/GenomicConsensus. Pacific Biosciences https://github.com/PacificBiosciences/GenomicConsensus (Accessed November 4, 2019).

2017a. rtg-tools: RTG Tools: Utilities for accurate VCF comparison and manipulation. Real Time Genomics https://github.com/RealTimeGenomics/rtg-tools (Accessed September 15, 2017).

2017b. vcflib: a simple C++ library for parsing and manipulating VCF files, + many command-line utilities. vcflib https://github.com/vcflib/vcflib (Accessed September 15, 2017).

